# Efficient Generation of Humanized Cloned Cattle via Embryonic Stem Cell-Mediated Dual-Round Site-Specific Editing

**DOI:** 10.64898/2025.12.20.695363

**Authors:** Zhiqiang Feng, Jinying Zhang, Shenao Tao, Peiru Lv, Xiaowei Zhang, Shunxin Wang, Minglei Zhi, He Zhang, Chang Zeng, Xin Li, Zengyuan Zhao, Yanhua Li, Tianzhi Chen, Yadi Teng, Yuhan Yang, Zimo Zhao, Hongkuan Ruan, Zhu Ma, Suying Cao, Shujing Li, Jianyong Han

**Affiliations:** State Key Laboratory of Animal Biotech Breeding, Frontiers Science Center for Molecular Design Breeding (MOE), College of Biological Sciences, China Agricultural University, Beijing, China; Henan Chuangyuan Biotechnology Co., Ltd; Animal Science and Technology College, Beijing University of Agriculture, Beijing, China; Shijiazhuang Tianquan Elite Dairy Co., Ltd. Shijiazhuang, Hebei 050200, People’s Republic of China; Beijing Dairy Cattle Center, Beijing 100192, People’s Republic of China

**Author notes:** These authors contributed equally: Zhiqiang Feng, Jinying Zhang, Shenao Tao, Peiru Lv. Correspondence should be addressed to: Suying Cao, Shujing Li, Jianyong Han.

## Abstract

Transgenic cloning allows genetic modification in cattle, yet its translational potential is constrained by low cloning efficiency and limited gene-editing capacity, due largely to the restricted proliferative ability of somatic donor cells. Pluripotent stem cells (PSCs), with their unlimited self-renewal ability, represent a promising alternative to overcome these bottlenecks. Here, we first established stable bovine epiblast stem cell lines (bEpiSCs) and achieved site-specific integration of the HCSN2-HLF-GFP cassette into the β-casein locus, with high transfection (72.07%) and correct integration efficiency (80.48%). Following Cre LoxP mediated excision of the selectable marker, we obtained marker free HCSN2-HLF bEpiSCs that retained pluripotency markers, tri lineage differentiation potential, and supported target protein expression in immortalized bovine mammary epithelial cells. When used as nuclear donors, these cells produced blastocysts with quality comparable to in vitro fertilization (IVF) embryos. Notably, cloning with HCSN2-HLF bEpiSCs resulted in the birth of twelve calves, with significantly higher birth and survival rates (30.71% and 16.74%, respectively) than those obtained using fibroblasts. Both PCR-based genotyping of the edited loci and microsatellite kinship analysis confirmed that all cloned calves were genetically derived from the HCSN2-HLF bEpiSCs. Collectively, this study establishes an efficient embryonic stem cell based platform for targeted gene editing and rapid production of multigene edited cloned cattle, offering a scalable strategy with strong potential for industrial translation.

## Dear Editor

Transgenic cloning enables genetic modification in livestock ^1–3^, with dairy cattle serving as effective bioreactors for the production of functional recombinant proteins in milk ^4^, thereby enhancing both biological activity and commercial value of dairy products. Humanized dairy models have been obtained to express key human proteins including lactoferrin ^5^, serum albumin ^6^, and lysozyme ^7^ in bovine milk, significantly improving its nutritional and functional properties. While translational potential of the current strategy is dual constrained by low cloning efficiency and limited gene editing capability due to the proliferative ability of donor somatic cells, which cannot tolerate continuous rounds of complex gene editing. Specifically, the survival rate of cloned animals typically remains as low as 1–5% ^5, 8^, which severely impedes the industrial-scale application of this technology. Endowed with inherent unlimited proliferative capacity, pluripotent stem cells tolerate continuous multiple rounds gene editing ^9^. This may effectively overcome the predicaments encountered in somatic cells. Here, we established a technical system to efficiently obtained cloned cattle from stable embryonic stem cells with at least three milk protein genes precise edits.

We selected human β-casein (*HCSN2*) and human lactoferrin (*HLF*), which are two key functional proteins in human milk, for transgene candidates. As a major nutritional structural protein, HCSN2 directly participates in regulating digestion, absorption, and mineral transport; while, HLF serves as a core immunodefensive protein and plays a central role in human milk-mediated immune protection. To perform dual-round site-specific editing, we use stable bovine epiblast stem cells (bEpiSCs) generated from early embryos for this study. We have designed a gene manipulation strategy (Fig. 1A) and successfully achieved site-specific integration of the HCSN2-HLF fragment into the third exon of the bovine β-casein gene by tandemly linking the *HCSN2* and *HLF* genes and adopting a two-step strategy assisted by the GFP fluorescent reporter system. Subsequently, the edited HCSN2-HLF bEpiSCs were used as donors for cell nuclear transfer.

**Fig. 1:**
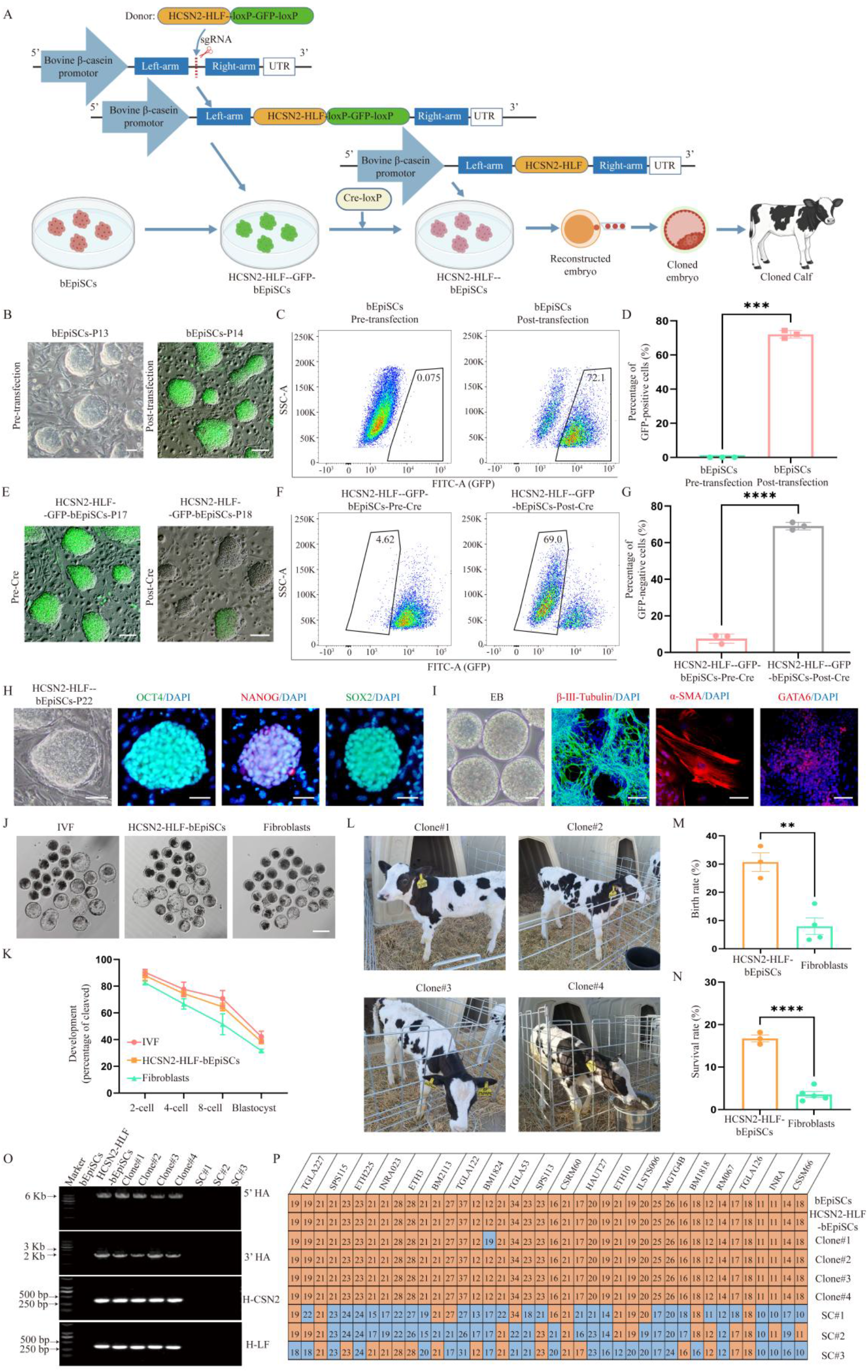
Generation of Cloned Cattle for Human β-Casein and Lactoferrin Production via bEpiSCs. A Schematic diagram of cloned cattle editing mediated by bEpiSCs. B Pre-transfection and post-transfection cellular status of bEpiSCs following transfection with the HCSN2-HLF-GFP plasmid. Scale bar, 50 μm. C The proportion of GFP-positive cells in bEpiSCs pre-transfection and post-transfection, with trapezoidal boxes indicating GFP-positive cells. D Bar plots showing the proportion of positive cells in bEpiSCs following transfection with the HCSN2-HLF-GFP plasmid as shown in C. Data are presented as mean ± SD, n = 3 replicates. \*\*\**P*<0.001. E Pre- and post-Cre transfection cellular status of HCSN2-HLF-GFP bEpiSCs. Scale bar, 50 μm. F The proportion of GFP-negative cells in HCSN2-HLF-GFP bEpiSCs pre- and post-Cre transfection, with trapezoidal boxes indicating GFP-negative cells. G Bar plots showing the proportion of negative cells in HCSN2-HLF-GFP bEpiSCs transfection with Cre as shown in F. Data are presented as mean ± SD, n = 3 replicates. \*\*\*\**P*<0.0001. H Cellular status and immunofluorescence staining of core pluripotency marker proteins OCT4 NANOG and SOX2 in HCSN2-HLF bEpiSCs. Scale bar, 50 μm. I *In vitro* embryoid body (EB) formation of HCSN2-HLF bEpiSCs and the capacity of EB for spontaneous differentiation into the three germ layers, along with immunofluorescence staining of the ectodermal neural-specific marker β-III-tubulin, mesodermal muscle-specific marker α-SMA, and endodermal-specific marker GATA6. Scale bar, 50 μm. J Schematic diagram of E7 blastocysts derived from *in vitro* fertilization (IVF), HCSN2-HLF bEpiSCs, and fibroblasts. Scale bar, 200 μm. K The line graph showing the early developmental capacity of different types of embryos. Data are presented as the mean ± SD., n = 3 replicates. L Healthy calves derived from HCSN2-HLF bEpiSCs via nuclear transfer technology, with the photograph taken on the 45th day of the calves’ survival. Clone#1 refers to cloned calf No.1, and so forth. M Bar plots showing a comparison of the birth rates of cloned calves derived from HCSN2-HLF bEpiSCs and fibroblasts. Data are presented as the mean ±SD., n = 3 replicates. \*\**P*<0.01. N Bar plots showing a comparison of the survival rates of cloned calves derived from HCSN2-HLF bEpiSCs and fibroblasts. Data are presented as the mean ±SD., n = 3 replicates. \*\*\*\**P*<0.0001. O Detection of the 5’ homology arm, 3’ homology arm, and target gene in cloned calves derived from HCSN2-HLF bEpiSCs, with bEpiSCs serving as the negative control and HCSN2-HLF bEpiSCs as the positive control. SC#1 denotes surrogate cow No.1, and so forth. Clone#1 was derived from SC#1; Clone#2 and Clone#3 were derived from SC#2; Clone#3 was derived from SC#3. P Calf parentage testing. Through microsatellite analysis, the allele fragment lengths at 20 loci were compared among bEpiSCs, HCSN2-HLF bEpiSCs, cloned calves, and surrogate cows; fragments consistent with those of bEpiSCs were labeled in orange, whereas inconsistent ones were labeled in blue.

We first successfully established three stable bEpiSC lines, which can be passaged for over 50 times without any signs of differentiation, from the epiblasts of three E7-stage blastocysts of high-yield dairy cows (Fig S1A). These cells consistently maintained a typical dome-shaped colony morphology (Fig S1B), positive alkaline phosphatase (AP) staining (Fig S1C), and a normal karyotype of 60 chromosomes (Fig S1D) during long-term passage. Immunofluorescence results demonstrated that the cells stably expressed the core pluripotency transcription factors such as OCT4, NANOG, and SOX2 (Fig S1E). Further sex identification was performed via specific PCR detection of the SRY gene, and the results showed that two lines were male and one was female (Fig S1F, G). The female cell line was selected as the candidate for subsequent gene editing in this study.

Initially, the HCSN2-HLF-GFP plasmid was transfected into bEpiSCs at passage 13 (Fig 1B). At 48 hours post-transfection, robust GFP fluorescence was observed in most colonies (Fig 1B). GFP-positive cells were then enriched via flow cytometry, achieving a sorting efficiency of 72.07% ± 2.25% (Fig 1C, D). The sorted GFP-positive cells were subjected to clonal expansion, and individual colonies were randomly selected for PCR-based genotyping (Fig. S2C). Amplification of fragments spanning the 5′ homology arm (5′HA: 6.1 kb), 3′ homology arm (3′HA: 4.1 kb), *HCSN2* (360 bp), and *HLF* (360 bp) confirmed that the proportion of correctly targeted colonies, termed as HCSN2-HLF-GFP bEpiSCs, was 80.48% ± 6.43% (Fig. S2E).

To ensure that cloned animals exclusively express the target protein and eliminate the potential risks associated with the reporter gene, site-specific excision was mediated by Cre recombinase transfection following the establishment of passage 17 HCSN2-HLF-GFP bEpiSCs (Fig. 1E). GFP-negative cell populations were isolated via flow cytometry, with an excision efficiency of 69.07% ± 2.00% (Fig. 1F, G). After clonal expansion of these GFP-negative cells, individual colonies were randomly selected for systematic genotyping, which included PCR validation of long homologous arms (5′HA: 6.1 kb; 3′HA: 2.1 kb) and target genes (*HCSN2*: 360 bp; *HLF*: 360 bp) (Fig. S2D). The results demonstrated that all tested colonies exhibited precise integration and stably harbored the intact *HCSN2* and *HLF* genes (Fig. S2F), culminating in the successful construction of a marker-free HCSN2-HLF bEpiSCs. Notably, the efficiency of large-fragment precise integration in stem cells is significantly higher than that in somatic cells, and the operation is convenient ^5, 10^.

Next, we analyzed the pluripotency characteristics of HCSN2-HLF bEpiSCs. We found that the cells retained canonical pluripotent morphological and molecular features after genetic editing. They maintained a dome-shaped colony morphology and positive AP staining (Fig. S3A), with sustained and stable nuclear expression of the core pluripotent transcription factors such as OCT4, SOX2, and NANOG (Fig. 1H, Fig. S3B). In addition, *in vitro* embryoid body (EB) formation assays showed that the cells were capable of self-assembling into EBs with intact structures and distinct outlines (Fig. S3C). Following spontaneous differentiation, cells derived from these EBs expressed markers specific to the three germ layers, such as ectodermal marker β-III-tubulin, mesodermal marker α-SMA, and endodermal marker GATA6, confirming their robust differentiation potential (Fig. 1I, Fig. S3D).

To validate the feasibility of the gene-editing strategy developed in this study, we first confirmed the suitability of immortalized bovine mammary epithelial cells (IBMECs) for subsequent functional assays via identification of mammary gland-specific markers (Fig. S4A). Subsequently, the editing strategy previously optimized in bovine embryonic stem cells (bEpiSCs) was applied in parallel to IBMECs. PCR amplification spanning homology arms and the target genes enabled the isolation of cell lines with site-specific integration of the HCSN2-HLF cassette (Fig. S4B). Furthermore, *in vitro* lactation induction assays combined with enzyme-linked immunosorbent assay (ELISA) analyses of both the supernatant and cell pellets revealed that endogenous bovine β-casein expression was significantly downregulated in edited cells, whereas the expression levels of exogenous human β-casein and human lactoferrin were markedly elevated (Fig. S4E, H, K). These findings demonstrate the efficacy and functionality of our gene-editing strategy at both the protein secretion and intracellular storage levels.

Subsequently, to systematically evaluate the developmental potential of HCSN2-HLF bEpiSCs as donor cells for nuclear transfer, we examined the early developmental efficiency of cloned embryos derived from this cell line. The results demonstrated that morphologically normal blastocysts could be successfully obtained using HCSN2-HLF bEpiSCs as donors (Fig. 1J). To further clarify the characteristics of their developmental capacity, we performed a horizontal comparison of the early embryonic developmental efficiency among the groups of *in vitro* fertilization (IVF), HCSN2-HLF bEpiSCs nuclear transfer, and fibroblast nuclear transfer. Interestingly, the early developmental capacity of HCSN2-HLF bEpiSCs-derived cloned embryos is significantly superior to that of the fibroblast cloning group and closer to that of IVF embryos (Fig. 1K, Table S1). Furthermore, cell counting analysis at the blastocyst stage showed that both the total cell number and the inner cell mass (ICM) cell number of blastocysts in the HCSN2-HLF bEpiSCs nuclear transfer group were significantly higher than those in the fibroblast group (Fig. S5A, B, C), indicating better embryonic developmental quality and further confirming that HCSN2-HLF bEpiSCs possess the potential to serve as high-quality donor cells for nuclear transfer.

Finally, 105 cloned embryos were generated using HCSN2-HLF bEpiSCs as nuclear donors and transplanted into 71 surrogate cows in three separate batches (Table S2), leading to the successful birth of 12 healthy cloned calves (Fig. 1L, Fig. S6A, Table S3). Quantitative analysis revealed that both the birth rate (30.71% ± 5.68%) and survival rate (16.74% ± 1.40%) in the HCSN2-HLF bEpiSCs group were significantly higher than those in the fibroblast group (birth rate: 7.14% ± 4.40%; survival rate: 3.91% ± 1.68%) (Fig. 1M, N, Table S2). Subsequent genotypic characterization of the 12 calves via cross-homologous arm PCR and target gene detection confirmed that all calves contained the expected integrated fragments (5′ HA: 6.1 kb; 3′ HA: 2.1 kb) along with intact *HCSN2* and *HLF* genes (Fig. 1O, Fig. S6B, C). Moreover, kinship analysis based on fragment length polymorphism of 20 short tandem repeats (STR) loci demonstrated that all cloned calves were genetically consistent with the parental bEpiSCs and HCSN2-HLF bEpiSCs, while showing no genetic association with the 11 surrogate cows (Fig. 1P, Fig. S6D, E, Fig. S7A, B, C). Collectively, these results confirm that all cloned calves were derived from HCSN2-HLF bEpiSCs, thereby validating the reliability and efficiency of this cell line as a nuclear transfer donor. Our study establishes an efficient and precise gene-editing cloning platform mediated by embryonic stem cells, enabling the rapid generation of multiplex gene-edited cloned cattle. It provides a scalable, integrated technological solution for large-scale pharmaceutical protein production via mammary gland bioreactors and the construction of humanized large-animal disease models.

## Notes

### Competing Interest Statement

The authors have declared no competing interest.

### Summary of Updates

The author's affiliation order has been corrected

### Funding

- the Biological Breeding-National Science and Technology Major Project (2023ZD0407503), the National Key R&D Program of China (2022YFD1302201), the Natural Science Foundation of China (32302746, 32370846), Hebei Province Science and Technology Plan (23227602Z), and Shijiazhuang City Science and Technology Project (221790262A), the 2115 Talent Development Program of China Agricultural University

