## Supplementary information for "Efficient Generation of Humanized Cloned Cattle via Embryonic Stem Cell-Mediated Dual-Round Site-Specific Editing"

[*]Correspondence should be addressed to:

### Materials and methods

#### Animal Handling and Ethical Statement

All mouse experiments and dairy cattle-related operations were approved in advance by the Animal Ethics Committee of China Agricultural University (AW10204202-3-1).

#### Preparation of Mouse Feeder Cells

For preparation of mouse feeder cells, embryos were isolated from E13.5 pregnant CD-1® (ICR) IGS mice. Following removal of visceral organs, heads, tails and limbs, trunk tissues were digested with 0.25% trypsin-EDTA (Gibco, 25300120) at 37 °C. The resulting cell suspension was seeded onto culture dishes containing MEF medium, which was composed of DMEM (Gibco, 11960044) supplemented with 10% (v/v) fetal bovine serum (FBS, Gibco, 16000044), 1% penicillin-streptomycin (Thermo Fisher Scientific, 15140122) and 1% GlutaMAX (Thermo Fisher Scientific, 35050061). Cells were cultured at 37 °C in a humidified incubator with 5% $CO_2$. Upon passage of primary cells to the third passage and achievement of confluence, cells were treated with 13.3 ng/mL mitomycin C for 3.5 hours to arrest proliferation. Cells were then washed with Dulbecco's phosphate-buffered saline (DPBS, Gibco, 21600044) and digested with 0.1% trypsin-EDTA to obtain feeder cells. After cell counting, feeder cells were aliquoted into cryovials at a density of $2.4\times10^6$ cells per vial and subjected to long-term storage via a programmed gradient cryopreservation method.

#### Isolation and Culture of Bovine Epiblast Stem Cells (bEpiSCs)

Isolation of the epiblast from high-yield dairy cow embryos and establishment of stable cell lines have been described in our previous study [1]. A brief description is as follows: Epiblast cells from high-yield dairy cow embryos were mechanically isolated and digested with TrypLE™ Express (Gibco, 12605010) for 3 minutes. The cells were then seeded onto mouse feeder cells containing 3i/LAF medium [1]. Subsequently, the cells were transferred to an incubator with 5% $O_2$, 5% $CO_2$ and saturated humidity for culture. After the clones grew for 2–3 days, they were digested with Accutase (Gibco, A1110501). When most clones detached, an equal volume of 3i/LAF medium was added to terminate digestion. The cells were then passaged at a ratio of 1:4, and this process was repeated every 2–3 days.

**Alkaline Phosphatase (AP) staining**

Alkaline Phosphatase (AP) staining was performed using the Alkaline Phosphatase Detection Kit (Millipore, SCR004). For the detailed AP staining procedure, reagent preparation, and precautions, refer to the instruction manual of the aforementioned kit.

**Immunofluorescence Staining**

After washing with DPBS, cells were fixed with 4% paraformaldehyde (PFA) at room temperature for 30 minutes. Following fixation, cells were rinsed with DPBS and then permeabilized with 0.01% Triton X-100 for 20 minutes. Subsequently, cells were washed with DPBS again and blocked with 3% BSA at room temperature for 1 hour. Primary antibodies were incubated at 4°C overnight, after which cells were washed three times with washing buffer (DPBS containing 0.1% Triton X-100 and 0.1% Tween 20). Secondary antibodies were then added and incubated at room temperature for 1 hour, followed by three washes with the same washing buffer. Finally, DAPI staining was performed to label cell nuclei, and images were observed and captured under a fluorescence microscope.

**Karyotype Analysis**

Prior to karyotype analysis, 1% KaryoMAX® Colcemid solution (Gibco, 15212012) was added to the culture medium of bEpiSCs and incubated for 2–3 hours to arrest cells at metaphase. Cells were then digested with TrypLE™ Express (Gibco, 12605010). Following centrifugation, approximately 50 μL of medium was retained to resuspend the cell pellet. Subsequently, 5 mL of pre-warmed 0.075 M KCl solution (Sigma, P5405) was slowly added, and hypotonic treatment was conducted at 37 °C for 30 minutes. Thereafter, 500 μL of freshly prepared fixative (3:1 v/v methanol:acetic acid) was added, and the mixture was gently inverted for thorough mixing. Cells were centrifuged at 1000 rpm for 5 minutes, and the supernatant was discarded. This fixation step was repeated twice with 5 mL of fixative each time, and the cell pellet was finally resuspended in 1 mL of fixative.

The cell suspension was dropped onto pre-cooled glass slides inclined at 45° from a height of 1 m, followed by air-drying at room temperature. Dried slides were stained with 10% Giemsa stain (Sangon, E6073140001) in the dark for 10–15 minutes and then dried in a 37 °C oven. Metaphase cells were observed and imaged under an oil immersion microscope, with more than 45 metaphase cells analyzed per sample.

**Embryoid body differentiation**

After enzymatic digestion, HCSN2-HLF-bEpiSCs were inoculated onto 35 mm low-adhesion culture dishes at a density of $1 \times 10^6$ cells per well. The cells were cultured in MEF growth medium for 5 to 7 days under shaking conditions—specifically, on a horizontal shaker set at 70 revolutions per minute (rpm). Thereafter, the formed Embryoid Bodies (EBs) were transferred to 12-well culture plates and maintained in the same growth medium for one week. During this period, the medium was refreshed every other day. Finally, the adherent cells obtained from this culture process were used for subsequent immunofluorescence staining experiments.

**Plasmid construction**

The PX330-Puro plasmid was constructed following the protocol described in previous literature [2]. For the exon region of the bovine CSN2 gene (Gene ID: 281099), four sgRNA sequences were designed using the CRISPOR online tool (http://crispor.tefor.net/) in accordance with the NGG PAM rule. These sgRNAs were synthesized by BGI Genomics, with their target sequences (20 bp flanking the PAM site) listed as follows:

sgRNA-1: 5′-aggctttccacaatctataa-3′ (SEQ ID No. 2),

sgRNA-2: 5′-ctggaagaactcaatgtacc-3′ (SEQ ID No. 3),

sgRNA-3: 5′-agtaacagtctctaatgatc-3′ (SEQ ID No. 4),

sgRNA-4: 5′-agaactcaatgtacctggtg-3′ (SEQ ID No. 5).

For constructing sgRNA expression plasmids, the PX330-Puro plasmid was first digested with BbsI restriction enzyme at 37 °C for 1 hour to generate the linearized vector. Each sgRNA was annealed to form double-stranded products, which were then individually ligated to the linearized vector using T4 DNA ligase. This ligation step ultimately yielded four recombinant plasmids: PX330-bCSN2 sgRNA-1-Puro, PX330-bCSN2 sgRNA-2-Puro, PX330-bCSN2 sgRNA-3-Puro, and PX330-bCSN2 sgRNA-4-Puro.

To assess the cleavage efficiency of each sgRNA, bEpiSCs at 70–80% confluence were seeded into 24-well plates and transfected using Lipofectamine 3000 (Thermo Fisher Scientific, L3000015). The transfection system per well contained 2 μg of one recombinant plasmid and 1 μL of P3000 reagent. These components were separately diluted in 50 μL of Opti-MEM, mixed gently, and incubated at room temperature for 15 minutes to form transfection complexes, which were subsequently added to the cells.

Post-transfection, quantitative insertion-deletion (indel) analysis of edited cells was performed via the TIDE (Tracking of Indels by DEcomposition) method. This approach resolves complex signals (e.g., double peaks or overlapping peaks) induced by heterozygous or biallelic editing by comparing Sanger sequencing electropherograms of edited and control cells, thereby enabling the identification and quantification of major indel types and their frequencies. Specifically, the sgRNA target sequence (20 nt upstream of the PAM) and sequencing files of control and edited cells were uploaded to the TIDE online platform (https://tide.nki.nl). The indel detection range was set to 0–50 bp, with all other parameters kept default. This analysis efficiently quantifies the mutation frequency in the edited cell population, providing a reliable quantitative basis for subsequent functional studies. The results indicated that sgRNA-4 exhibited the highest cleavage efficiency and was thus selected for subsequent experiments (Fig. S2A, B).

The construction method of the gbCSN2-Donor-hCSN2-hLF-loxp-eGFP-Puro-loxp plasmid (abbreviated as HCSN2-HLF-GFP) is described as follows. The complete nucleotide sequence of this plasmid is provided in SEQ ID No. 1. Using the Donor-OCT4-A5-3×flag-PNP-tdtomato vector as the backbone, we designed the left and right homology arms at 5.5 kb upstream and 2.0 kb downstream of the target sgRNA site, respectively. First, the backbone vector and the PCR-amplified 5′ homology arm fragment were double-digested with MluI and AgeI. After recovering the linearized vector and the homology arm fragment, T4 ligase was used for ligation to complete the insertion of the 5′ homology arm. Subsequently, following the same strategy, the aforementioned intermediate vector and the 3′ homology arm fragment were digested and ligated with ClaI and PacI, thereby completing the assembly of the homology arms on both sides. Finally, the key functional element synthesized by BGI Genomics—a fragment containing hCSN2-hLF-LOXP-EF-1α-EGFP-P2A-PuroR-SV40 poly(A)-LOXP—was integrated into the donor vector with assembled homology arms via homologous recombination to obtain the complete HCSN2-HLF-GFP plasmid.

The construction protocol for the pCAG-Cre-IRES-Puro plasmid is detailed below: Using the pCAG-IRES-Puro vector (complete nucleotide sequence provided in SEQ ID No. 6) as the backbone, its original small fragment was replaced with the coding sequence (CDS) of Cre recombinase (Gene ID: 2777477) via the EcoRI and AgeI restriction enzyme sites. The remaining regions of the vector were retained unchanged, resulting in the generation of the pCAG-Cre-IRES-Puro expression plasmid.

**Transfection and Screening of bEpiSCs**

Transfection and screening of bEpiSCs were conducted as follows: First, bEpiSCs were harvested via digestion with Accutase (Gibco, A11105-01). For each transfection reaction, $1\times10^6$ cells were subjected to co-transfection with 2 μg of PX330-bCSN2 sgRNA-4-Puro plasmid and 2 μg of HCSN2-HLF-GFP donor vector using the Neon NxT Electroporation System. The electroporation parameters were set as 1200 V, 20 ms, and 2 pulses, with the aim of site-specifically integrating the hCSN2-hLF-loxp-eGFP-Puro-loxp sequence into the third exon of the BCSN2 gene.

After the bEpiSCs to be transfected grow stably, GFP-positive cells were sorted using a MoFlo XDP flow cytometer (Beckman Coulter) with excitation at 488 nm and a 710/50 bandpass filter (primary screening based on the GFP reporter system). The sorted cells were then seeded into 48-well plates for monoclonal expansion. To verify the gene editing efficiency, genomic DNA was directly extracted from monoclonal cell lines using a cell lysis buffer (Invitrogen, AM8723) and utilized as the template for PCR amplification. Ultimately, the HCSN2-HLF-GFP bEpiSCs cell line was successfully isolated through the detection of cross-homologous arms and the target gene.

Following the acquisition of the stable HCSN2-HLF-GFP bEpiSCs cell line, 2 μg of pCAG-Cre-IRES-Puro plasmid was electroporated into this cell line to induce Cre-loxP-mediated excision for the removal of the eGFP reporter gene. Subsequently, GFP-negative cells were collected via flow sorting. After expansion, detection of cross-homologous arms and the target gene was performed to obtain the HCSN2-HLF-bEpiSCs cell line. This step effectively employed the Cre-loxP system to efficiently eliminate the eGFP marker, ultimately generating a bEpiSCs line harboring only the HCSN2 and HLF genes.

**Differentiation of the HCSN2-HLF-bEpiSCs**

HCSN2-HLF-bEpiSCs were initially cultured in 3i/LAF medium for 48 hours, followed by medium replacement with BM differentiation medium supplemented with BMP4 (10 ng/mL), SB-431542 (5μM), and FGF2 (10 ng/mL). Upon gradual transformation of cell morphology to a fibroblast-like phenotype, the cells were subcultured and continuously maintained in the differentiation medium for 1 week until they displayed a homogeneous, long spindle-shaped, fibroblast-like appearance.

**Extraction of Genomic DNA**

Genomic DNA from cell and tissue samples was extracted using the TIANamp Genomic DNA Kit (TIANGEN, DP304).

**Preparation of Dairy Cow Ear Marginal Fibroblasts**

Ear marginal tissue samples were harvested from healthy dairy cows at a commercial dairy farm. Upon collection, tissues were subjected to a stepwise sterile disinfection protocol: initially rinsed with 75% (v/v) ethanol for 30 s, followed by three consecutive washes with DPBS supplemented with 1% (v/v) penicillin-streptomycin (double antibodies). Excess hair was meticulously removed using sterile forceps, and cartilaginous components were excised with a sterile surgical blade to isolate homogeneous dermal tissue fragments. The trimmed tissues were transferred to 1.5 mL sterile centrifuge tubes, minced into 1–2 $mm^3$ pieces, and homogenized with 200 L FBS to form a uniform tissue slurry. The slurry was evenly plated onto the bottom of T25 cell culture flasks pre-loaded with mouse embryonic fibroblast (MEF) culture medium. Flasks were then placed in a humidified incubator maintained at 37 °C with 5% (v/v) $CO_2$ for primary culture, allowing the outgrowth of dairy cow ear marginal fibroblasts.

**In vitro maturation of oocytes**

Bovine ovaries were collected from a slaughterhouse, preserved in normal saline at 30 °C, and transported to the laboratory within 2 hours. Upon arrival, the ovaries were rinsed three times with normal saline pre-warmed to 38 °C. Follicles with a diameter of 2–8 mm were aspirated using a 10 mL syringe fitted with an 18-gauge needle to harvest cumulus-oocyte complexes (COCs). The COCs were placed in maturation medium for culture at 38.5 °C in an incubator with 5% $CO_2$ and saturated humidity for 16–17 hours. The formulation of the maturation medium can be found in previous reports [3]. Subsequently, excess cumulus cells were removed by treatment with 0.1% hyaluronidase, and mature oocytes that had extruded the first polar body were selected for subsequent experiments.

**Cloned Embryo Production**

he enucleation of oocytes was performed in M199 medium (Gibco, 11150-059) supplemented with 10% FBS. After enucleation, donor cells were microinjected into the cytoplasm of oocytes, and the operating medium was M199 containing 2% FBS. Cell fusion was achieved using an electrofusion instrument (CFB16-HB) with the parameters set as 22 V and 10 μs. Reconstructed embryos were treated with 5 μM ionomycin (Sigma, D2629) for 5 minutes, followed by transfer to culture medium containing 2 mM 6-DMAP (Sigma, I3909) for activation over 4 hours. Activated embryos were cultured in IVC medium (Bioscience, BOIVC2501) at 38.5°C in a 5% $CO_2$ incubator. The cleavage rate of embryos was recorded 48 hours post-culture, and the blastocyst development rate was counted on day 7 of culture.

**In vitro fertilization (IVF) of oocytes**

Bovine frozen semen was retrieved from the liquid nitrogen tank and thawed in a 38.5°C water bath. After thawing, the semen was transferred into a 15 mL centrifuge tube containing 6 mL of semen washing medium, centrifuged at 1800 rpm for 5 minutes, the supernatant was discarded, and the washing process was repeated once. Subsequently, semen washing medium was added to resuspend the spermatozoa; the sperm density was adjusted to $2\times10^6$ cells/mL under a microscope using a hemocytometer, and the suspension was placed in an incubator for standing and subsequent use.

The formulation of BO basal medium was as follows: 112 mM NaCl (Sigma-Aldrich, S5886), 4.02 mM KCl (Sigma-Aldrich, P5405), 2.25 mM $CaCl_2 \cdot 2H_2O$ (Sigma-Aldrich, C5670), 0.83 mM $NaH_2PO_4 \cdot H_2O$ (Sigma-Aldrich, S9638), 0.52 mM $MgCl_2 \cdot 6H_2O$ (Sigma-Aldrich, M2393), 37 mM $NaHCO_3$ (Sigma-Aldrich, S5761), 1.25 mM sodium pyruvate (Sigma-Aldrich, P5280), and 10 μg/mL heparin (Sigma-Aldrich, H3149). On this basis, semen washing medium was supplemented with 10 mM caffeine (Sigma-Aldrich, C8960) and 4 mg/mL bovine serum albumin (BSA, Sigma-Aldrich, A1470), while fertilization medium was only supplemented with 4 mg/mL BSA.

COCs that had undergone in vitro maturation culture for 22–24 hours were washed three times with fertilization medium, then transferred into 50 μL fertilization medium droplets that had been equilibrated in an incubator for at least 2 hours, with 15–20 COCs placed in each droplet. Fifty microliters of diluted sperm suspension was added to each fertilization droplet, and sperm-oocyte co-incubation was performed in an incubator at 38.5°C, 5% $CO_2$, and saturated humidity for 8–18 hours.

### Synchronization treatment of recipient cows

Healthy recipient cows in optimal physiological condition were selected and subjected to estrus synchronization treatment following this protocol: an intravaginal progesterone insert (Intravaginal Progesterone Insert, DEC International, CIDR 1380) was intravaginally placed on Day 0; 0.6 mg of Cloprostenol Sodium Injection (Qilu Animal Health Products Co., Ltd., Beiduofu) was intramuscularly administered on Day 7; the progesterone insert was removed on Day 10; on Day 19, morphologically normal and well-developed blastocysts were selected and transferred into the uterus of recipient cows that had undergone estrus synchronization treatment, followed by embryo transfer implementation.

### Enzyme-Linked Immunosorbent Assay（ELISA）

Immortalized bovine mammary epithelial cells (IBMECs) were purchased from Shanghai Yaji Biotechnology Co., Ltd (YS152820C). These cells were genetically engineered using the same gene editing strategy as that for bEpiSCs, thereby enabling them to harbor the HCSN2 and HLF genes. After the edited mammary epithelial cells were passaged and had adhered to the culture vessel, the medium was replaced with induction medium (composed of 10% FBS, 1% antibiotics-antimycotic, 89% DMEM medium, and 1 $\mu g \cdot mL^{-1}$ prolactin (Cloud-clone corp. Wuhan, RPA846Bo01)) for continuous incubation for 48 hours. Upon completion of induction, the cell supernatant and cell pellet were collected separately. ELISA kits were utilized to quantify the levels of bovine β-casein, human β-casein, and human lactoferrin in both the cells and supernatants, with all specific procedures conducted strictly in accordance with the manufacturers' instructions. The kits employed included the Human β-Casein (β-CN) ELISA Research Kit (MM-63257H1), Bovine β-Casein (CSN2; β-CN) ELISA Research Kit (MM-50471O1), and Human Lactoferrin (LF/LTF) ELISA Research Kit (MM-1248H1), all of which were procured from Jiangsu Meimian Industrial Co., Ltd. To verify the specificity of the kits, three types of kits were respectively used to detect the corresponding protein contents in human milk and cow's milk. The results showed that the bovine β-casein detection kit could only recognize β-casein in cow's milk, with no signal detected in human milk samples (Fig S4D). Similarly, the human β-casein detection kit and the human lactoferrin detection kit only produced specific responses to the target proteins in human milk, and no corresponding proteins were detected in cow's milk (Fig S4G, J). The above results indicated that each kit exhibited good species specificity.

**Microsatellite Detection and Analysis**

Umbilical cord tissues of newborn calves and uterine tissues of recipient cows were harvested for genomic DNA isolation. Thereafter, 20 bovine microsatellite loci were assayed utilizing the GTyping® Bovine Microsatellite Marker (STR/SSR) Detection Kit (Glbizzia) in conjunction with quantitative real-time PCR (qPCR) technology. The assayed loci were classified into two categories: Category I encompasses 12 SSR loci recommended by the International Society for Animal Genetics (ISAG) for routine kinship verification, including TGLA227, BM2113, TGLA53, ETH10, SPS115, TGLA126, TGLA122, INRA23, ETH3, ETH225, BM1824, and BM1818; Category II comprises 8 SSR loci jointly recommended by ISAG and the Food and Agriculture Organization of the United Nations (FAO) for livestock genetic studies, namely SPS113, CRM060, HAUT27, ILSTS006, MGTG4B, RM067, INRA063, and CSSM66.

**Primer sequences**

Bovine SRY (501bp): sex identification of bEpiSCs.

F: ATGTTCAGAGTATTGAACGAC, R: TGAGTATGTGGTCTTGGCACA.

5’HA (6.1Kb): detection of the 5'-cross-homologous arm in HCSN2-HLF-bEpiSCs, HCSN2-HLF-bEpiSC, HCSN2-HLF-IBMECs and calves.

F: ATGCTGAAGCTGAAACTCCTATACTTTAG, R: CAAATTGGGAAAGGAGTA-
-CATCAAGGAAT.

3’HA-1 (4.1Kb): detection of the 3'-cross-homologous arm in HCSN2-HLF-bEpiSCs.

F: ATGGAGAGCGACGAGAG, R: TTGTCAAAGTTTTTATTTCTTGCACTG.

3’HA-2 (2.1Kb): detection of the 3'-cross-homologous arm in HCSN2-HLF-bEpiSC, HCSN2-HLF-IBMECs and calves.

HCSN2 (360bp): Detection of target genes in HCSN2-HLF-bEpiSCs, HCSN2-HLF-bEpiSCs , HCSN2-HLF-IBMECs and calves.

F: GCAGAGTGATGCCTGTCCTT, R: ACAGCTCTCTGAGGGTAGGG.

HLF(360bp): Detection of target genes in HCSN2-HLF-bEpiSCs, HCSN2-HLF-bEpiSCs , HCSN2-HLF-IBMECs and calves.

F: CCTGCTCTTCAACCAGACGG, GCCATGGCAAGATGGCAG.

**Statistical analysis**

Figure 1D, G, K, M, N, Table S1, and Figure S4 D, E, G, H, J, K were subjected to Welch's t-test analysis, n = 3 replicates. Figure S2B, Figure S5B, C were analyzed using the two-way ANOVA multiple comparison test, with n = 3 replicates. Data presented in Table S2 are derived from the following sources: Fibroblasts 1 group was generated in the present study; Fibroblasts 2, 3, and 4 groups were sourced from previously published literature. Consequently, the "Fibroblasts" group in Figure 1M, N is aggregated from the combined data of Fibroblasts 1–4 groups.

**Drawing Software**

Figure 1A and Figure S1A were created with BioGDP.com.

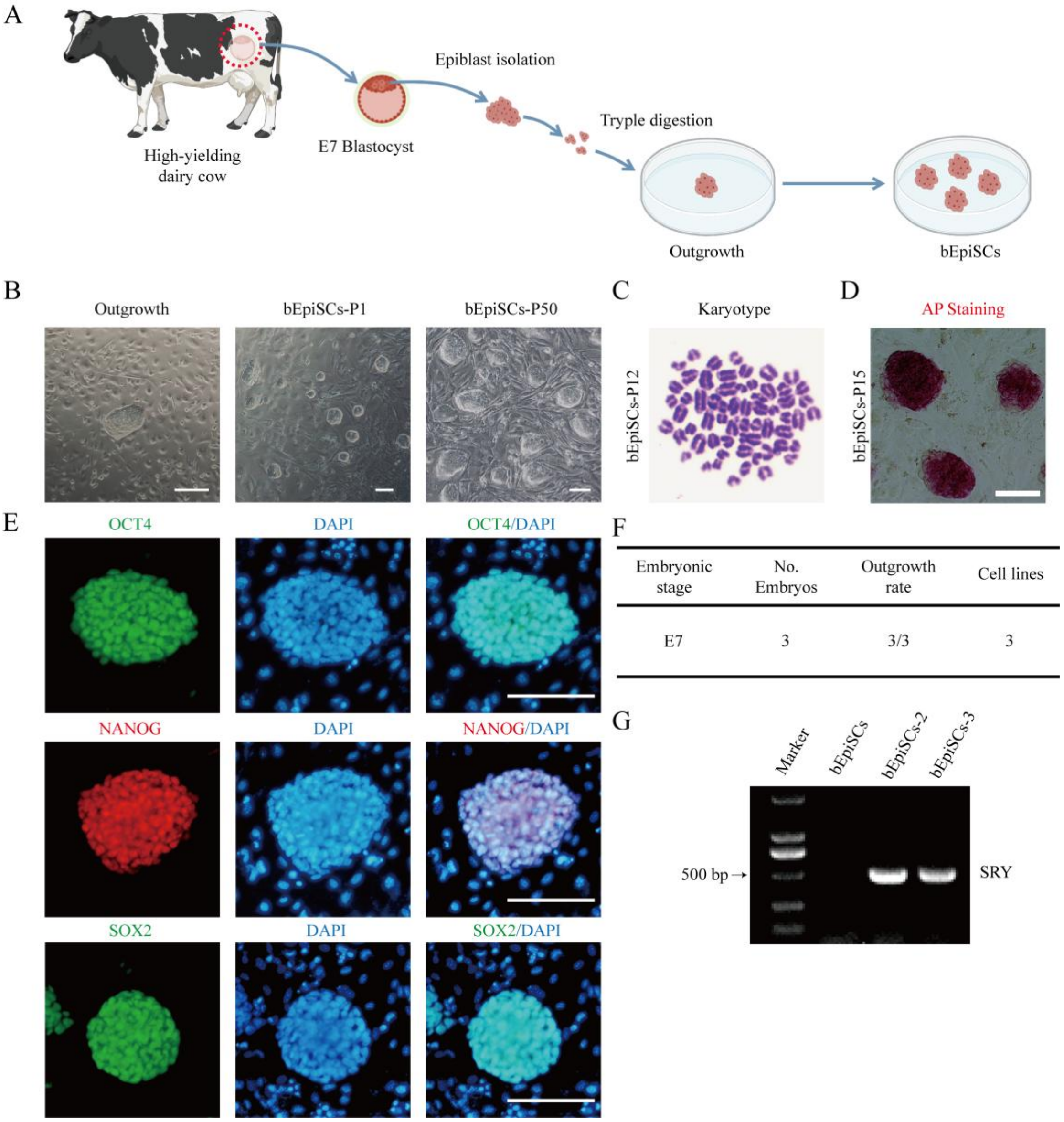


**Fig. S1: Establishment and Identification of bEpiSCs from High-Yield Dairy Cows**

A Schematic diagram of cell line establishment from E7 blastocysts of high-yield dairy cows.

B Schematic diagram of outgrowth derived from EPI, passage 1 (P1) bEpiSCs, and passage 50 (P50) bEpiSCs. Scale bar, 100 μm.

C Karyotype analysis of bEpiSCs, with karyotypes of at least 45 metaphase cells counted for each cell line

D Alkaline phosphatase staining analysis of bEpiSCs. Scale bar, 100 μm.

E Immunofluorescence staining of core pluripotency marker proteins OCT4, NANOG and SOX2 in bEpiSCs. Scale bar, 100 μm.

F Statistical analysis of cell line establishment from E7 blastocysts of high-yield dairy cows.

G Sex identification of bEpiSCs: the presence of an SRY band indicates males, while the absence indicates females.

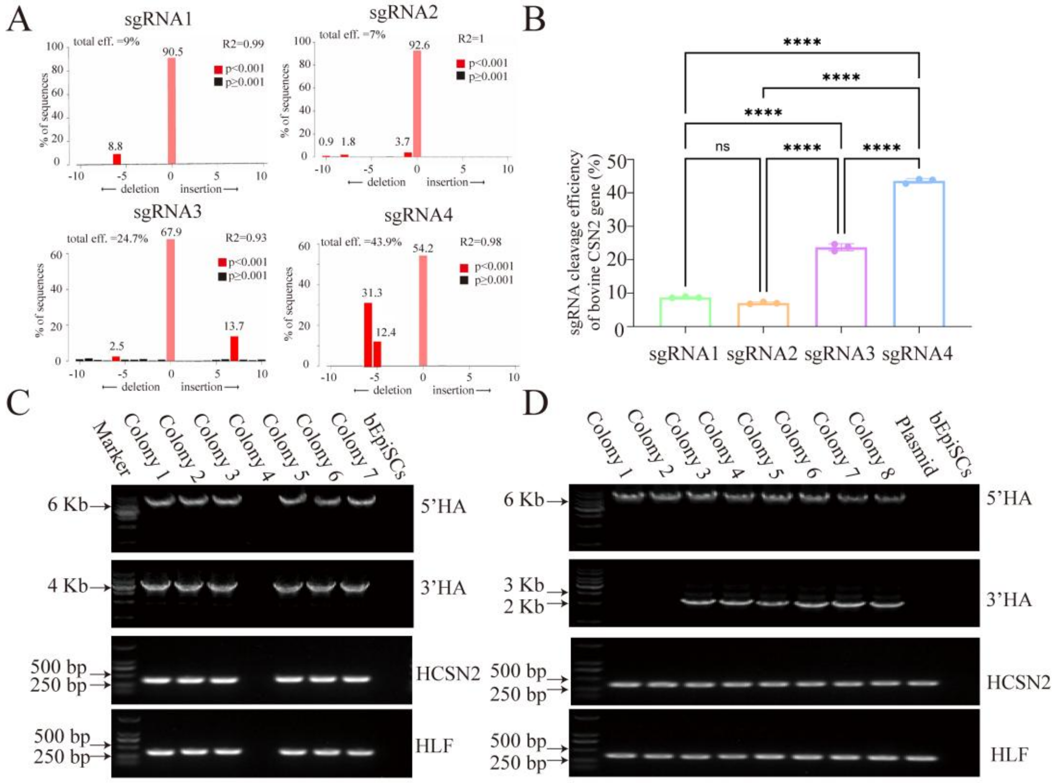


E

| Cell lines | No. of colonies | 5'HA (6.1Kb) | 3'HA (4.1Kb) | HCSN2 (360 bp) | HLF (360 bp) | No. of positive colonies (%±SD%) |
|---|---|---|---|---|---|---|
| bEpiSCs | 5 | 0 | 0 | 0 | 0 | 0 |
| HCSN2-HLF--GFP-bEpiSCs | 41 | 33 | 33 | 33 | 33 | 33 (80.48±6.43) |

F

| Cell lines | No. of colonies | 5'HA (6.1Kb) | 3'HA (2.1Kb) | HCSN2 (360 bp) | HLF (360 bp) | No. of positive colonies (%±SD%) |
|---|---|---|---|---|---|---|
| HCSN2-HLF--GFP-bEpiSCs | 10 | 10 | 0 | 10 | 10 | 0 |
| HCSN2-HLF--bEpiSCs | 40 | 40 | 40 | 40 | 40 | 100% |

**Fig. S2: Identification of HCSN2-HLF-GFP bEpiSCs and HCSN2-HLF bEpiSCs.**

A Analysis of sgRNA cleavage efficiency.

B Bar plot comparison of the cleavage efficiencies of sgRNA 1, sgRNA 2, sgRNA 3, and sgRNA 4 on the bovine CSN2 gene. Data are presented as the mean ± SD, n = 3 replicates. ****$P<0.0001$.

C Detection of the 5' homology arm, 3' homology arm and target gene in HCSN2-HLF-GFP bEpiSCs: GFP-positive clones were randomly selected for detection (Colony 1–7), with bEpiSCs as the negative control, n = 5 replicates.

D Detection of the 5' homology arm, 3' homology arm and target gene in HCSN2-HLF bEpiSCs: GFP-positive clones (colony 1–2) and GFP-negative clones (colony 3–8) were randomly selected for detection, with bEpiSCs as the negative control and HCSN2-HLF-GFP Plasmid as the positive control for the target gene, n = 5 replicates.

E Statistics of positive clones in HCSN2-HLF-GFP bEpiSCs and bEpiSCs. Data are presented as the mean ± SD, n = 5 replicates.

F Statistics of positive clones in HCSN2-HLF-GFP bEpiSCs and HCSN2-HLF bEpiSCs. n = 5 replicates.

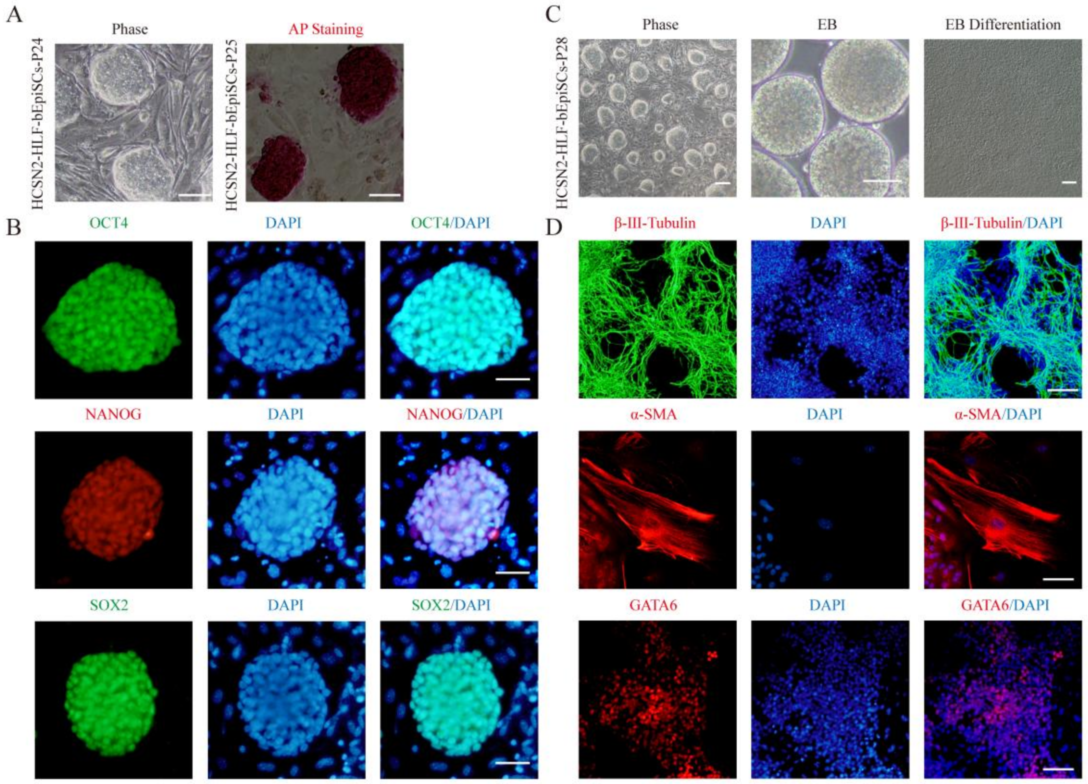


Fig. S3: Pluripotency Characterization of HCSN2-HLF-GFP bEpiSCs. Related to Fig. 1H, I.

A Bright-field images of HCSN2-HLF-GFP bEpiSCs at passage 24 and alkaline phosphatase (AP) staining of the cells at passage 25. Scale bar, 100 μm.

B Immunofluorescence staining of core pluripotency marker proteins OCT4, NANOG and SOX2 in HCSN2-HLF-GFP bEpiSCs. Scale bar, 50 μm.

C HCSN2-HLF-GFP bEpiSCs at passage 28 retained the capacity to form EB and undergo spontaneous differentiation in vitro. Scale bar, 100 μm.

D EB can differentiate randomly into cells of the three germ layers. Immunofluorescence staining was performed for the ectodermal neural-specific marker β-III-tubulin, mesodermal muscle-specific marker α-SMA, and endodermal-specific marker GATA6. Scale bar, 50 μm.

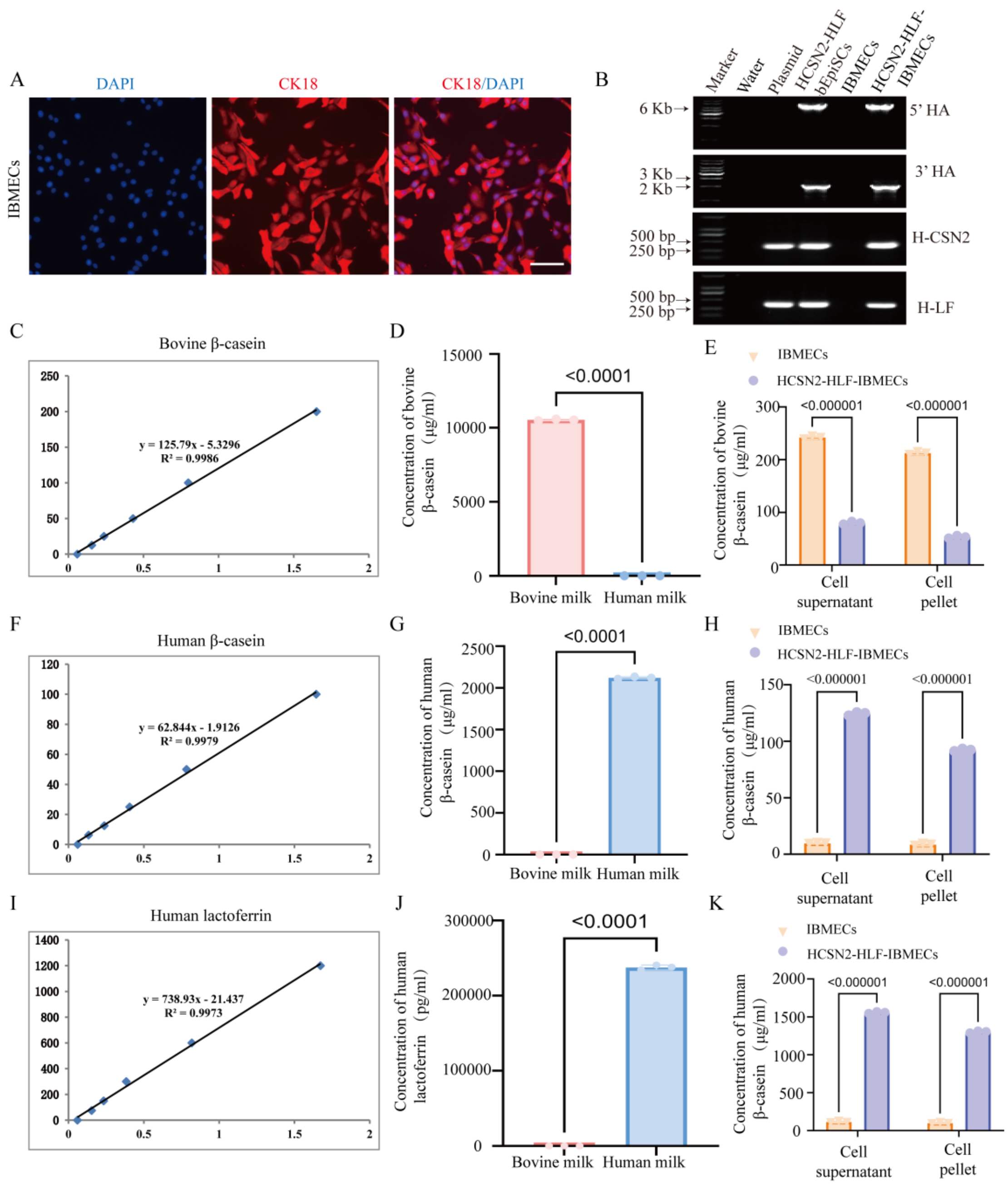


**Fig. S3: Pluripotency Characterization of HCSN2-HLF-GFP bEpiSCs. Related to Fig. 1H, I.**

A Bright-field images of HCSN2-HLF-GFP bEpiSCs at passage 24 and alkaline phosphatase (AP) staining of the cells at passage 25. Scale bar, 100 μm.

B Immunofluorescence staining of core pluripotency marker proteins OCT4, NANOG and SOX2 in HCSN2-HLF-GFP bEpiSCs. Scale bar, 50 μm.

C HCSN2-HLF-GFP bEpiSCs at passage 28 retained the capacity to form EB and undergo spontaneous differentiation in vitro. Scale bar, 100 μm.

D EB can differentiate randomly into cells of the three germ layers. Immunofluorescence staining was performed for the ectodermal neural-specific marker β-III-tubulin, mesodermal muscle-specific marker α-SMA, and endodermal-specific marker GATA6. Scale bar, 50 μm.

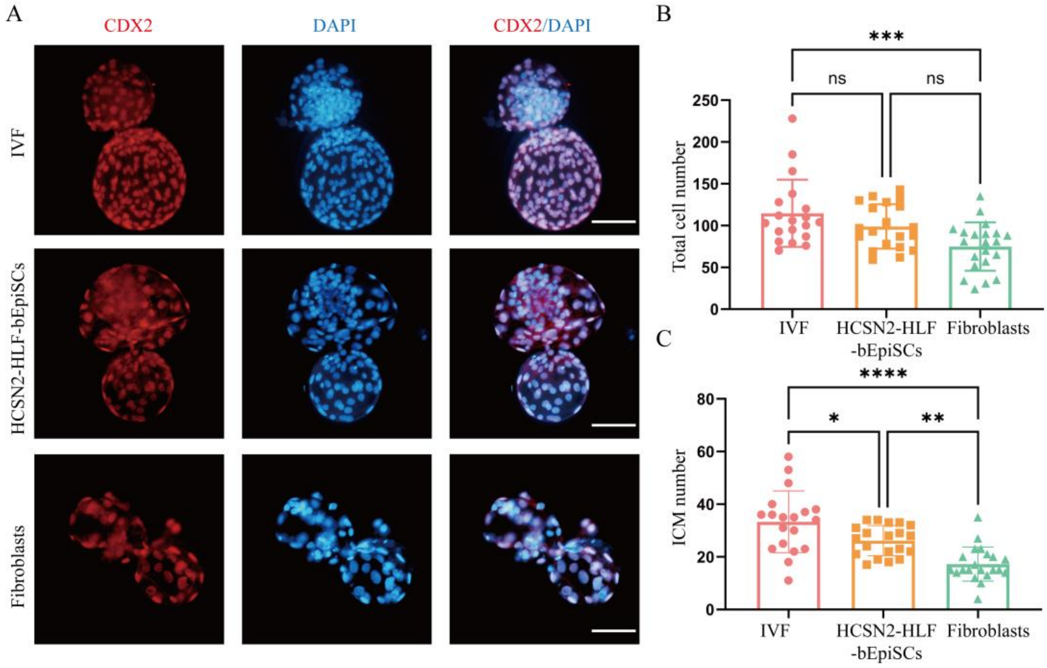


**Fig. S5: Evaluation of the quality of different types of blastocysts. Related to Fig. 1J.**

A Immunofluorescence staining of three types of blastocysts derived from IVF, HCSN2-HLF bEpiSCs, and fibroblasts: CDX2 was used to label trophectoderm cells, and DAPI was used to label the total number of cells. Scale bar, 100 μm.

B Bar plot showing the statistics of total cell numbers in blastocysts from three groups: IVF, HCSN2-HLF bEpiSCs, and fibroblasts. Data are presented as mean ± SD (IVF, n = 22; HCSN2-HLF bEpiSCs, n = 22; fibroblasts, n = 21).

C Bar plot showing the statistics of inner cell mass (ICM) counts in blastocysts from three groups: IVF, HCSN2-HLF bEpiSCs, and fibroblasts. The ICM count was calculated as total cell number minus trophectoderm cell number. Data are presented as mean ± SD (IVF, n = 22; HCSN2-HLF bEpiSCs, n = 22; fibroblasts, n = 21).

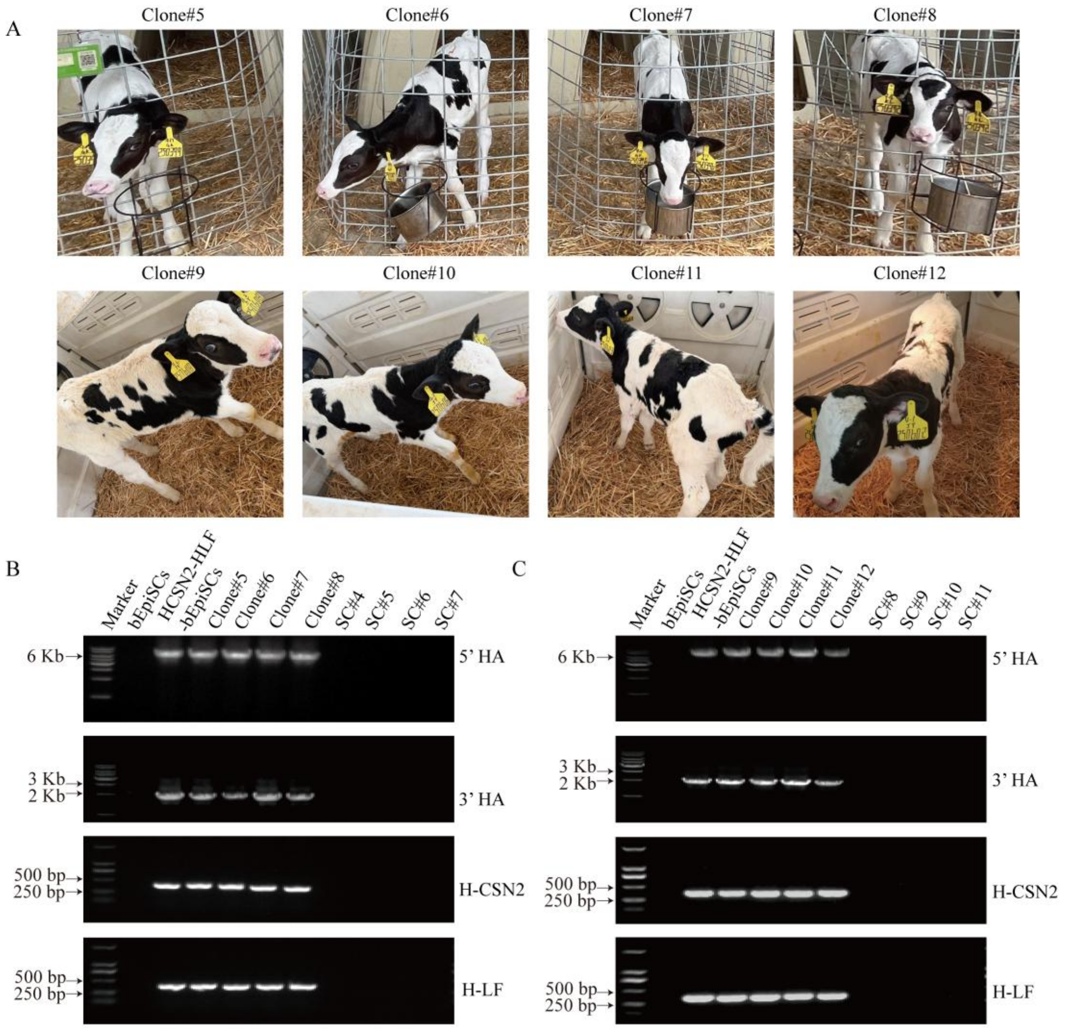


D

| | TGLA227 | | SPS115 | | ETH225 | | INRA023 | | ETH3 | | BM2113 | | TGLA122 | | BM1824 | | TGLA53 | | SPS113 | | CSRM60 | | HAUT27 | | ETH10 | | ILSTS006 | | MGTG4B | | BM1818 | | RM067 | | TGLA126 | | INRA | | CSSM66 | |
|---|---|---|---|---|---|---|---|---|---|---|---|---|---|---|---|---|---|---|---|---|---|---|---|---|---|---|---|---|---|---|---|---|---|---|---|---|---|---|---|---|
| bEpiSCs | 19 | 19 | 21 | 21 | 23 | 23 | 21 | 21 | 28 | 28 | 21 | 21 | 27 | 37 | 12 | 12 | 21 | 34 | 23 | 23 | 16 | 21 | 17 | 20 | 19 | 21 | 19 | 20 | 25 | 26 | 16 | 18 | 12 | 14 | 17 | 18 | 11 | 11 | 14 | 18 |
| HCSN2-HLF -bEpiSCs | 19 | 19 | 21 | 21 | 23 | 23 | 21 | 21 | 28 | 28 | 21 | 21 | 27 | 37 | 12 | 12 | 21 | 34 | 23 | 23 | 16 | 21 | 17 | 20 | 19 | 21 | 19 | 20 | 25 | 26 | 16 | 18 | 12 | 14 | 17 | 18 | 11 | 11 | 14 | 18 |
| Clone#5 | 19 | 19 | 21 | 21 | 23 | 23 | 21 | 21 | 28 | 28 | 21 | 21 | 27 | 37 | 12 | 12 | 21 | 34 | 23 | 23 | 16 | 21 | 17 | 20 | 19 | 21 | 19 | 20 | 25 | 26 | 16 | 18 | 12 | 14 | 17 | 18 | 11 | 11 | 14 | 18 |
| Clone#6 | 19 | 19 | 21 | 21 | 23 | 23 | 21 | 21 | 28 | 28 | 21 | 21 | 27 | 37 | 12 | 12 | 21 | 34 | 23 | 23 | 16 | 21 | 17 | 20 | 19 | 21 | 19 | 20 | 25 | 26 | 16 | 18 | 12 | 14 | 17 | 18 | 11 | 11 | 14 | 18 |
| Clone#7 | 19 | 19 | 21 | 21 | 23 | 23 | 21 | 21 | 28 | 28 | 21 | 21 | 27 | 37 | 12 | 12 | 21 | 34 | 23 | 23 | 16 | 21 | 17 | 20 | 19 | 21 | 19 | 20 | 25 | 26 | 16 | 18 | 12 | 14 | 17 | 18 | 11 | 11 | 14 | 18 |
| Clone#8 | 19 | 19 | 21 | 21 | 23 | 23 | 21 | 21 | 28 | 28 | 21 | 21 | 27 | 37 | 12 | 19 | 21 | 34 | 23 | 23 | 16 | 21 | 17 | 20 | 19 | 21 | 19 | 20 | 25 | 26 | 16 | 18 | 12 | 14 | 17 | 18 | 11 | 11 | 14 | 18 |
| SC#4 | 17 | 18 | 21 | 21 | 24 | 24 | 17 | 21 | 22 | 22 | 21 | 21 | 17 | 20 | 12 | 13 | 21 | 23 | 19 | 24 | 16 | 20 | 20 | 20 | 14 | 18 | 17 | 20 | 17 | 20 | 16 | 18 | 12 | 12 | 17 | 18 | 10 | 11 | 17 | 11 |
| SC#5 | 19 | 22 | 23 | 23 | 24 | 24 | 15 | 17 | 22 | 27 | 15 | 21 | 17 | 21 | 17 | 17 | 22 | 23 | 23 | 24 | 20 | 21 | 20 | 23 | 19 | 21 | 19 | 19 | 17 | 20 | 17 | 18 | 12 | 14 | 18 | 18 | 10 | 11 | 19 | 19 |
| SC#6 | 19 | 22 | 21 | 21 | 23 | 24 | 15 | 17 | 28 | 28 | 15 | 19 | 27 | 31 | 12 | 17 | 22 | 30 | 23 | 23 | 16 | 21 | 17 | 23 | 18 | 22 | 20 | 20 | 17 | 22 | 17 | 18 | 12 | 14 | 18 | 18 | 10 | 11 | 18 | 18 |
| SC#7 | 18 | 18 | 21 | 21 | 23 | 24 | 21 | 21 | 28 | 28 | 15 | 19 | 26 | 37 | 12 | 17 | 30 | 34 | 23 | 23 | 23 | 21 | 17 | 20 | 14 | 22 | 17 | 20 | 21 | 24 | 18 | 20 | 12 | 14 | 17 | 18 | 11 | 11 | 19 | 19 |

E

| | TGLA227 | | SPS115 | | ETH225 | | INRA023 | | ETH3 | | BM2113 | | TGLA122 | | BM1824 | | TGLA53 | | SPS113 | | CSRM60 | | HAUT27 | | ETH10 | | ILSTS006 | | MGTG4B | | BM1818 | | RM067 | | TGLA126 | | INRA | | CSSM66 | |
|---|---|---|---|---|---|---|---|---|---|---|---|---|---|---|---|---|---|---|---|---|---|---|---|---|---|---|---|---|---|---|---|---|---|---|---|---|---|---|---|---|
| bEpiSCs | 19 | 19 | 21 | 21 | 23 | 23 | 21 | 21 | 28 | 28 | 21 | 21 | 27 | 37 | 12 | 12 | 21 | 34 | 23 | 23 | 16 | 21 | 17 | 20 | 19 | 21 | 19 | 20 | 25 | 26 | 16 | 18 | 12 | 14 | 17 | 18 | 11 | 11 | 14 | 11 |
| HCSN2-HLF -bEpiSCs | 19 | 19 | 21 | 21 | 23 | 23 | 21 | 21 | 28 | 28 | 21 | 21 | 27 | 37 | 12 | 12 | 21 | 34 | 23 | 23 | 16 | 21 | 17 | 20 | 19 | 21 | 19 | 20 | 25 | 26 | 16 | 18 | 12 | 14 | 17 | 18 | 11 | 11 | 14 | 11 |
| Clone#9 | 19 | 19 | 21 | 21 | 23 | 23 | 21 | 21 | 28 | 28 | 21 | 21 | 27 | 37 | 12 | 12 | 21 | 34 | 23 | 23 | 16 | 21 | 17 | 20 | 19 | 21 | 19 | 20 | 25 | 26 | 16 | 18 | 12 | 14 | 17 | 18 | 11 | 11 | 14 | 18 |
| Clone#10 | 19 | 19 | 21 | 21 | 23 | 23 | 21 | 21 | 28 | 28 | 21 | 21 | 27 | 37 | 12 | 12 | 21 | 34 | 23 | 23 | 16 | 21 | 17 | 20 | 19 | 21 | 19 | 20 | 25 | 26 | 16 | 18 | 12 | 14 | 17 | 18 | 11 | 11 | 14 | 18 |
| Clone#11 | 19 | 19 | 21 | 21 | 23 | 23 | 21 | 21 | 28 | 28 | 21 | 21 | 27 | 37 | 12 | 12 | 21 | 34 | 23 | 23 | 16 | 21 | 17 | 20 | 19 | 21 | 19 | 20 | 25 | 26 | 16 | 18 | 12 | 14 | 17 | 18 | 11 | 11 | 14 | 18 |
| Clone#12 | 19 | 19 | 21 | 21 | 23 | 23 | 21 | 21 | 28 | 28 | 21 | 21 | 27 | 37 | 12 | 12 | 21 | 34 | 23 | 23 | 16 | 21 | 17 | 20 | 19 | 21 | 19 | 20 | 25 | 26 | 16 | 18 | 12 | 14 | 17 | 18 | 11 | 11 | 14 | 18 |
| SC#8 | 20 | 22 | 21 | 23 | 24 | 24 | 17 | 19 | 22 | 22 | 15 | 21 | 21 | 27 | 12 | 13 | 34 | 35 | 21 | 23 | 21 | 21 | 17 | 23 | 19 | 21 | 17 | 19 | 17 | 20 | 16 | 19 | 12 | 12 | 17 | 18 | 10 | 10 | 17 | 19 |
| SC#9 | 19 | 19 | 21 | 21 | 22 | 23 | 15 | 21 | 22 | 22 | 14 | 15 | 17 | 26 | 12 | 17 | 21 | 23 | 23 | 24 | 16 | 16 | 17 | 21 | 19 | 22 | 17 | 19 | 20 | 26 | 16 | 18 | 12 | 12 | 18 | 18 | 11 | 11 | 17 | 19 |
| SC#10 | 18 | 18 | 21 | 23 | 24 | 24 | 19 | 21 | 23 | 28 | 15 | 19 | 17 | 20 | 12 | 17 | 26 | 35 | 23 | 24 | 16 | 16 | 17 | 17 | 21 | 22 | 17 | 21 | 20 | 20 | 16 | 18 | 12 | 13 | 18 | 21 | 10 | 10 | 19 | 19 |
| SC#11 | 19 | 22 | 21 | 23 | 19 | 23 | 17 | 21 | 28 | 28 | 20 | 21 | 27 | 37 | 12 | 17 | 23 | 35 | 21 | 21 | 16 | 18 | 17 | 23 | 18 | 19 | 19 | 20 | 20 | 22 | 17 | 18 | 12 | 12 | 18 | 18 | 10 | 10 | 19 | 19 |

**Fig. S6: Identification of cloned calves derived from HCSN2-HLF bEpiSCs. Related to table S4, 5.**

A Images of Clone#5–8 at 45 days postnatal. Images of Clone#9–12 at 20 days postnatal. Clone #5 follows the numbering sequence of Clone #4 in Fig. 1K, representing the 5th cloned calf; Clone #6 represents the 6th cloned calf, and so on.

B Detection of the 5' homology arm, 3' homology arm, and target gene in Clone#5–8. bEpiSCs were used as the negative control, and HCSN2-HLF bEpiSCs as the positive control. SC#4 is the abbreviation for the 4th surrogate cow, SC#5 for the 5th surrogate cow, and so on. Clone#5 was derived from surrogate cow SC#4, Clone#6 from SC#5, Clone#7 from SC#6, and Clone#8 from SC#7.

C Detection of the 5' homology arm, 3' homology arm, and target gene in Clone#9–12. bEpiSCs were used as the negative control, and HCSN2-HLF bEpiSCs as the positive control. SC#8 is the abbreviation for the 8th surrogate cow, SC#9 for the 9th surrogate cow, and so on. Clone#9 was derived from surrogate cow SC#8, Clone#10 from SC#9, Clone#11 from SC#10, and Clone#12 from SC#11.

D Parentage testing of Clone#5–8 calves. Through microsatellite analysis, the allele fragment lengths at 20 loci were compared among bEpiSCs, HCSN2-HLF bEpiSCs, cloned calves, and surrogate cows; fragments consistent with those of bEpiSCs were labeled in orange, whereas inconsistent ones were labeled in blue.

E Parentage testing of Clone#9–12 calves.

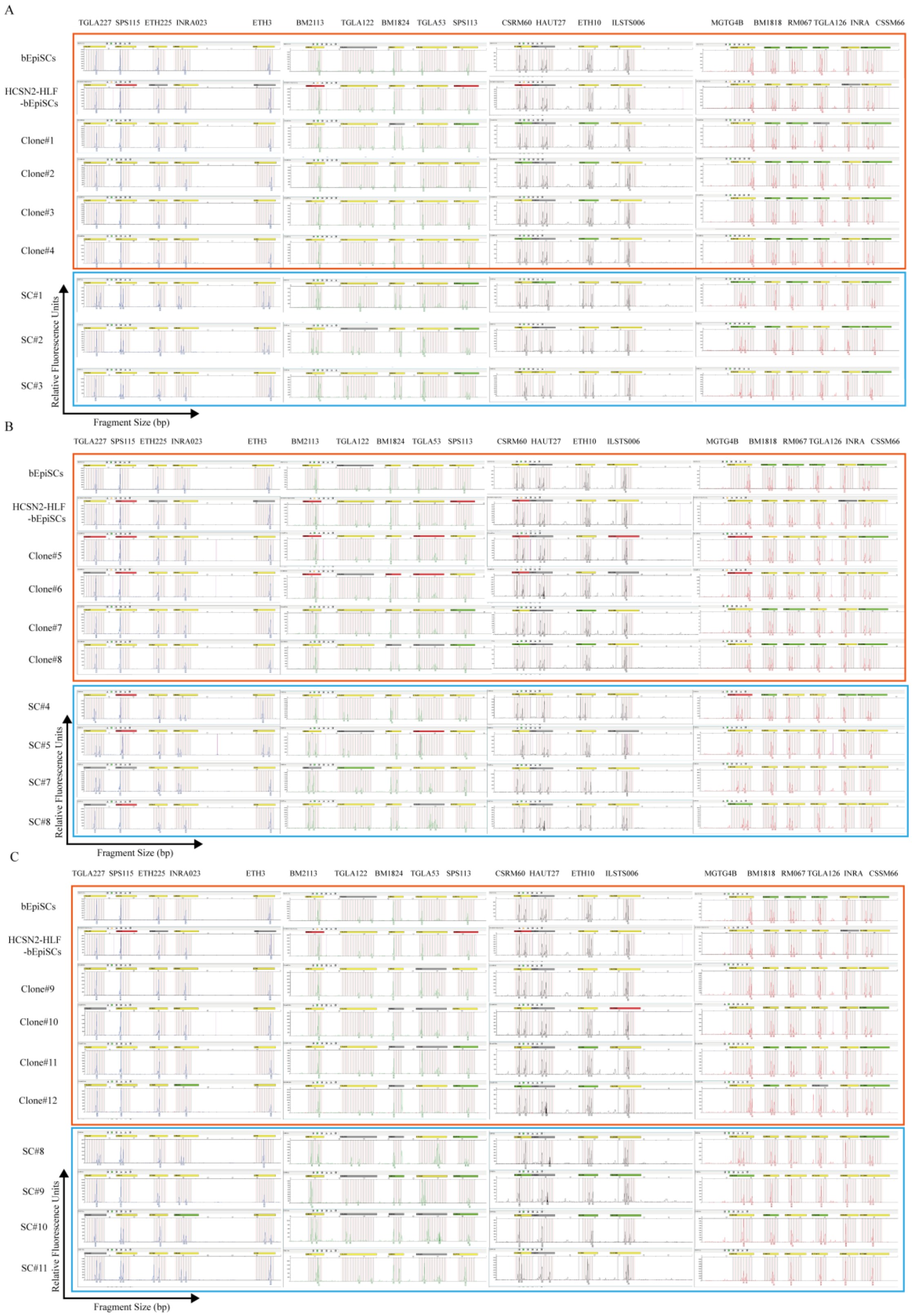


**Fig. S7: Parentage Identification of Cloned Calves Derived from HCSN2-HLF bEpiSCs. Related to Fig. 1O, Fig. S6D, E.**

A Microsatellite identification results of bEpiSCs, HCSN2-HLF bEpiSCs, Clone#1–4, and SC#1–3: the abscissa represents the allele fragment length, and the ordinate represents the fluorescence intensity of the samples.

B Microsatellite identification results of bEpiSCs, HCSN2-HLF bEpiSCs, Clone#5-8, and SC#4-7.

C Microsatellite identification results of bEpiSCs, HCSN2-HLF bEpiSCs, Clone#9-12, and SC#8-11.

**Table S1 Developmental Rates of Different Embryo Types**

| Group | No. of cultured | No. of 2-cell (%±SD%) | No. of 4-cell (%±SD%) | No. of 8-cell (%±SD%) | No. of blastocyst (%±SD%) |
|---|---|---|---|---|---|
| IVF | 166 | 150 (90.39±1.93) | 130 (77.72±4.76) | 118 (70.72±5.32) | 70 (42.17±3.71) |
| HCSN2-HLF-bEpiSCs | 149 | 131 (87.91±3.12) | 111 (74.49±2.10) | 96 (64.50±2.56) * | 57(38.42±3.39)* |
| Fibroblast | 120 | 99 (82.51±2.13) *** | 80 (66.72±3.45) ** | 62 (51.52±6.96) ** | 38 (31.67±4.12) *** |

* *P* <0.05, ** *P* <0.01, *** *P* <0.001 as compared with IVF group, by Welch's t *t*-test, n=3 replicates.

No. of 2-cell (%±SD%): The proportion of embryos with the number of blastomeres ≥ 2 relative to the total number of cultured embryos.

No. of 4-cell (%±SD%): The proportion of embryos with the number of blastomeres ≥ 4 relative to the total number of cultured embryos.

No. of 8-cell (%±SD%): The proportion of embryos with the number of blastomeres = 8 relative to the total number of cultured embryos.

No. of blastocyst (%±SD%): The proportion of blastocysts relative to the total number of cultured embryos.

**Table S2 In Vivo Development of Cloned Embryos Derived from Different Cell Types**

| Cell line | Replicate | No. Blastocyst | Surrogates | Pregnant (%) | Birth (%) | Survival (%) |
|---|---|---|---|---|---|---|
| HCSN2-HLF-bEpiSCs | 1 | 30 | 24 | 12 (40.00) | 6 (25.00) | 4 (16.67) |
| | 2 | 30 | 22 | 12 (75.00) | 8 (36.37) | 4 (18.18) |
| | 3 | 45 | 25 | 16 (59.25) | 8 (30.77) | 4 (15.38) |
| Fibroblasts 1 | 1 | 40 | 20 | 14 (70.00) | 3 (12.00) | 1 (5.00) |
| | 2 | 60 | 30 | 15 (50.00) | 4 (13.33) | 2 (6.67) |
| Fibroblasts 2 [4] | 1 | 462 | 129 | 28 (21.70) | 11 (8.52) | 5 (3.87) |
| Fibroblasts 3 [5] | 1 | 97 | 31 | 14 (45.16) | 1 (3.22) | 1 (3.22) |
| Fibroblasts 4 [6] | 2 | 143 | 49 | 9 (18.37) | 2 (4.08) | 1 (2.04) |
| | 2 | 119 | 37 | 5 (13.51) | 1 (2.70) | 1 (2.70) |

Pregnant (%): The proportion of cows with 30-day gestation relative to the total number of surrogate cows.

Birth (%): The proportion of live-born cloned calves relative to the total number of surrogate cows

Survival (%): The proportion of cloned calves surviving to the age of one month relative to the total number of surrogate cows.

Fibroblasts 1 were generated in this study

Fibroblasts 2, 3, and 4 groups were sourced from previously published literature.

**Table S3 Transplantation and Birth Weight of Transgenic Cloned Calf**

| Surrogate Cow | No. embryos transferred (ET) per recipient | Cloned Calf | Calf Ear Tag Number | Birth Weight (Kg) |
|---|---|---|---|---|
| SC#1 | 2 | Clone#1 | 250489 | 46 |
| SC#2 | 2 | Clone#2 | 250483 | 38.5 |
| | | Clone#3 | 250484 | 54 |
| SC#3 | 2 | Clone#4 | 250485 | 51 |
| SC#4 | 2 | Clone#5 | 250399 | 45.5 |
| SC#5 | 1 | Clone#6 | 250397 | 40.5 |
| SC#6 | 1 | Clone#7 | 250396 | 51 |
| SC#7 | 1 | Clone#8 | 250392 | 33 |
| SC#8 | 2 | Clone#9 | 250608 | 45 |
| SC#9 | 1 | Clone#10 | 250606 | 72 |
| SC#10 | 2 | Clone#11 | 250603 | 36 |
| SC#11 | 1 | Clone#12 | 250602 | 53 |

**SEQ ID No.1:**

AAAAATAAACAAATAGGGGTTCCGCGCACATTTCCCCGAAAAGTGCCACCTGACGTCT
AAGAAACCATTATTATCATGACATTAACCTATAAAAATAGGCGTATCACGAGGCCCTTTC
CGTTGTAAAACGACGGCCAGTCGAACCACGCGATGCGAACACACGCGTATTCTCTGGT
TCTTGATTTATTTTATAAATTAATCTTAATAGTTATACTTCACATAGATAGGAATTTATTATA
TTTGGATAATCTCATGGAAAGGATTAAATACTCCATCTATATGAGTAATGCTGAACTATCT
ACTCCTACCTAATAATTTGTCAGAATTCACTAATTCTGTGTTATATTGTTTCTAAATCTGA
ATCATTATATGAATCCTCAGTATTTTATTTTCCTGCCTCTATATTTTGGAATTTATTAAACA
GTGCTTCAAATAATTTTTAGGAAACAAGTTTTTCATAACAGCTTTATCTCTAAATAGCTT
TAGTATCTTGAAAAAGTAATACAAATTCTCAAATCCTCAGTTTCCTCTTCTCTAAATATAT
TAAAAATATTCTATGGATGATATCTCTTAATATTTATTTTTTTGGCCATCCAGCACAGCTT
ATGGGATCTTAGTTCCCCAGTGAGGGATTATACCCATGGCCACTGCAGTGAAAGTACAA
AATCCTAAACTGAACTCACCAGAGATTTCCCTATATCTCCTCTAGTTCTTTTTTCTGAAT
ATTTTTGGTCCCTTTATTGTACTCTTCATCCAACTTTTCTATTGCTTTCTTTCTTAAAGATA
TCATTTGCATGGTTTCAGCTAGAATATATCTGAATCTCAGGACTGCATATTCAGATTCATC
GGCCAATATGGAAAAAAACTTTGGCTGAACAGATCATGCTTATAAAAATTAGTACTAGA
ACATCTTGCTTTGACTATATCTTGCTCCTCACCCAGGGTTATCTAATAAAATTTCCCCATA
TGAAATTCTTTTGCCTTATAAAAATGACAACTCTTAGGTAACATTGCAAAAATTTGAGT
TGCCCATATGGCCCTTTGGCTTCTTACTGGTCATTGTGTTCTGAGGCTTACCTGGACAGC
GGTACCTGATGTCATCTTCAAATGCTGGCTTTTTAATTTTCCCTTGGACAAGCTTCTTTC
TTTAGTATATTGTTAAGGATTTCTTTGATCAAGATTTTACCTACTTTTCTGGTCCAAATCC
AATCCATCTCTATCAATTAATGTAATTCAAAATTGGTGAGAGACAGTCATTAGGAAATTC
TCTGTTCAATGCACAATATGTAAAGCATCTTCCTGAGAAAAGGGAAATGTTGAATGGG
AAGGACATGCTTTCTTTTGTATTTCTTTTCTCAGAAATCACACTTTTTTGCCTGTGGCCT
TGGCAACCAAAAGCTAACACATAAAGAAAGGCATATGAAGTAGCCAAGGCCTTTTCTA
GTTATATCTATGACACTGAGTTCATTCATCATTTATTTTCCTGACTTCCTCCTGGGTCCAT
ATGAGCAGTCTTAGAATGAATATTAGCTGAATAATCCAAATGCATAGTAGATGTTGATTT
GGGTTTTCTAAGCAATACAAGACTTCTATGACAGTGAGATGTATTACCATCCAACACAC
ATCTCAGCATGATATAAATGTAAGGTATATTGTGAAGAAAAATTATCAATTATGTCAAAG
TGCTTACTTTAGAAGATCATCTATCTGTTCCAAAGCTGTGAATATATATATTGAACATAAT

TAATAGACGAAACAAACCTTGTAAAAATGAGTAGTGTAAAATACAACTACATTTATGAA
CATCTATCACTAAAGAGGCAAAGAAAGTTGAGGACTGCTTTTGTAAATGGGCTCTTATT
AATGAAAAGTACTTTTGAGGTCTGGCTTAGACTCTATTGTAGTACTTATGGTAAGACCC
TCCTCTTGTCTGGGCTTTTCATTTTCTTTCTTCCTTCCCTCATTTGCCCTTCCATGAATAC
TAGCTGATAAACATTGACTCACTATAAAAGATATGAGGCCAAACTTGAGCTGTCCCATT
TTAATAAATCTGTATAAATAATATTTGTTCTACAGAAGTATTCTCTAAATAAATGTTACTT
TCTCTCTTAAAATCCCTCAACAAATCCCCACTATCTAGAGAATAAGATTGACATTCCCTG
GAGTCACAGCATGCTTTGTCTGCCATTATCTGACCCCTTTCTCTTTCTCTCTTCTCACCT
CCATCTACTCCTTTTTCCTTGCGATACATGACCCAGATTCACTGTTTGATTTGGCTTGCA
TGTGTGTGTGCTGAGTTGTGTCTCACTCTTGTCAACCCCATGAATGACAGTCCACCAG
GCTCCACTATTCCCAGTTTAGAATACTGGAGTGGATTGTGTTTCCTACTTCATTTGATTAT
TTTAGTGACTTTTAAATTTTTTTCCATATTCAGGAGGCTATTCTTTCCCTTTTAGTCTATA
CTGTCTTCGCTCTTCAGGTTCTAAGCTATCATCATGTGCTTGTTAGCTTGTTTCTTTCTCC
ATTATAGCATAAACACTAACAACTATTTCAGGTTAGCATGAGATTGTGTTCTTTGTGTGG
CCTGTGTATTTCTGGTGTGTATTAGAATTTACCCCATGATCTCTAAAGACCCACCGAATA
CTAAAGAGACCTCATTTGTAGTTACAATAATTTGGGGACTGGGCCAAAACTTCCGTGTG
TCCCAGCCAAGGTCTGTAGCTACTGGACAATTTAAATTCCTTTATCAGATTGTGAATTAT
TCCCTTTTAAATGCTCCCCAGAATTTTTGGGGACAGAAAAATAGGAAGAATTCATTTTC
TAATCATGCAGATTTCTAGGAATTTCAAATCCACTATTGGTTTTATTTCAAGCCACAAAA
TTAGCATGCCATTAAATACTATATATAAACAACCACAAAATCAGATCATTTATCCATTCAG
CTCCTCCTTCACTTCTTGTCCACTACTTTGGAAAAAAGGTAAGAATCTCAGATATAATTT
TCATTGTATCTGCTACTCATCTTTATTTTCAGACTAGGTTAAAATGTAGAAAGAACATAA
TTGCTTAAAATAGATCTTAAAAATAAGGATGTTTAAGATAAGGTTTACAGTATTTTCAGC
AAATTTGTTAAAATAGAAGCAACTATAAAGATTTGTAACAGTGGTTGCTATTTTCTTTAC
CACGAGACTAGTTTACAGGCTGTATTAAAAGATCTTTTCTTGAAATTAATATTTTCAATT
TGATTTAAACATACCTCAGCCATAAAGGCAAGCACATTTTATTTATACTATGGGAATTTG
AATAATTGTTACTGAAGAAGCTCTACCACAAAAAAGTTTATAGAGCTAGCATATTTAGT
CAGAGAGATAAGAGGGTTGTTAGGATACATGTGCTATTTGAAAGGTATTTATAAAAAGA
AGAGTATATTTATTAAAATTGCTCAGAACATCCAAATTTCAGGTTTATCATTTATCTTACA
ATATTTTCCAAAAATATTAAAATAGATACATGAAATACAGAAGTAAATTAAAGAGAAAG

TATTTTATTTTGTAAAAAAAATTCTAGGTTGGACAGGGAGTACCAGGAAACAAAAAAC
AATGAAAAATGTGATCTGACAGAAATTATAGCTCAAAGTATAGTAGTCAGTAATGAAAT
GGCTTAAAAATTGGCATATAAAATGCTAATTATAAAATAAACAAAATGTAATAATACCCT
CCCTACATGTAATGAACTCTGAGTATTATACTCTTTTTTGAAGTCTTGACAATGAAAATT
TATTTAGACTTTTATAGACATCTTGGATAAAGTTAAACAAATTACGAATTAGCATCCATG
AGAAAAATATAGAAAAAATTTCTTAATGTAGTTTGCAATCTGGGATTGAAGATGTGTGT
CAAGAGATGTTGATGGCGAGAACATTTTTTTTTCAAGAACTTATAAAAATGCGACAAA
ACAAACCATTTTATACATTTTGGTCAAAAATAGTATGTATTTTATTTTATGCTACAAGGAG
AAGTAGTCTAAAGTGAGGACTGGGCAAGAGAATCTGACACCCTGGTAAGTCACCGAG
AGATAAGTACACAGTTCTCTGTAGAGAAAATAAGCATAGTGTATGATCTCTAAAATTGT
GTGGGACAAAGGGGAGATAACATTAGGCATGTGGGGATGAAGACTGAGTACAGAAGA
AACAATCTAGTCAGTCCAAGAAAACATGTGGATCAATGGGACAAATAGAAGAAATGCT
AAAGTGAAGCAGAAGTCTTACTGGAAATAAAAGATATGAGGAAGACAAACATTCATGA
AAATCACTTAGTTTAGTAGAGAAAAGATAAAAATAAAGTATTACCTTCTTCTTCATATAC
ATTGTTTGATCAGATGCCCCTCAGTAAAACTGAGTCTCCAACAGAACTGAAACTTTTAT
ATTTTGTTTCACTGCTCTAATCCCAGAATCTAAGACATATCTGGCGATAAAAAATTAATA
AATAAATATTTTTAATAAGTTAAATCAATCACTTAATTTTTCTGTAAGTATCTGTGACTTC
TCTTCTGTCTTTCCAAAAAACACTCATAAGTACTGTGAATAAGATGAAAAGAGTGAAAT
AAGATATAGGCTGTTAGCTGAAAACATCTGGATGGCTGGGAGTGAAGCATTAACTTGA
AATGTAAGATTAATGAGTAATAGTAAATTTTAACCTTGGCCGTATGATAAAATGTCTATTA
ATATTTTTCTAAAATACAGGGCTTTTTGTTTTTGCCATGAGGTTTGCAGGATCTTGGTTC
CCTGATGAGGGATCAAACCTGGGCTCCCCTGGGAGCACGGAGTCTTGAGATTTTGTAT
TATACACTATCTTTGGTTTCTTTTAAAGGGAAGTAATTCTACTTAAATAAGAAAATAGAT
TGACAAGTAATACACTATTTCCACATCTTCCCATTCCCAGGAATTGAGAGCCATGAAGG
TCCTCATCCTTGCCTGCCTGGTGGCTCTGGCCCTTGCAAGAGAGGTTAAATACAGAAA
AAATGTTGAAATAAATAAGACTAGTACTATCTGCCTATGTGTAGAAAATCGCCATTACCA
ACATTGTAAATGTATAAATAATGCGCAATCTCAGATTTTTTTTGAATGCTAAGAAAGTCA
TTTACGTTCTATCCACTATCTCAGTAGTATCCTATGGGACCACAAGTCTGAGTCTAGTGC
TTTCTATAGTATTGTACCATCTGTACCATCAATCCCTAAAGAAAAAAGAAAATAAACCA
ATAAGCAACAGACTAACAAGAAGGAACACAGATAAGAACAAAAAGTGAGTAATATTG

696 CATAAATACAATTGCATGCATATACAATCTAGATAAATATATCTTATTCCAGTGATGAAAT

697 ATTTGTATCCCTTACTGTAGAGTGCTAGGTTTAGCTATGTCTATTCAACACAGGATGATA

698 CTCCAGAGGATGGTATATCAGACAACAATAATAAATATGTTGATAATTATAATAAAAAGT

699 GTTTCAGTAAAAATTAAAATAACTCCCTTTCTGTTACCCATAAAAACTCTTCATTAAAGT

700 TAAACAAAAATATACTAATGAAAGTTACTAAATTTTAAAGACTCTCGAAAGACATATAA

701 CATTTTTATTTTTCAGATTTGTGAAATAGATAGCTCTGAATAAAGCAAGTAAAAATTAGG

702 TAGGAAAATATTTTATAATGAGTTGACTGTGGGAACTAAAGTGTTTTTTTTTCTCTTTAG

703 CTGGAAGAACTCACCGGTGGATCCGGGCTCTGGTGCAACCAATTTCTCTCTTCTTAAA

704 CAAGCCGGTGATGTGGAGGAGAACCCCGGACCCGGATCCATGAAGGTCCTCATCCTCG

705 CCTGCCTGGTGGCTCTTGCTCTTGCAAGGGAGACCATAGAAAGCCTTTCAAGCAGTGA

706 GGAATCTATTACAGAATACAAGAAAGTTGAGAAGGTTAAACATGAGGACCAGCAGCA

707 AGGAGAGGATGAACACCAGGATAAAATCTACCCCTCTTTCCAGCCACAGCCTCTGATC

708 TATCCATTCGTTGAACCTATCCCCTATGGTTTTCTTCCACAAAACATTCTGCCTCTTGCT

709 CAGCCTGCTGTGGTGCTGCCTGTCCCTCAGCCTGAAATAATGGGAGTCCCTAAAGCTA

710 AAGACACTGTCTACACTAAGGGCAGAGTGATGCCTGTGCTTTAAATCTCCAACGATAC

711 CCTTTTTTGACCCTCAAATCCCAAAACTCACTGATCTTGGAAATCTGCATCTTCCTCTG

712 CCTCTGCTCCAGCCCTTGATGCAGCAGGTCCCTCAGCCTATTGCTCAGACTCTTGCCCT

713 TCCCCCTCAGCCCCTGTGGTCTGTTCCCTCAGCCCAAAGTCCTGCCTATCCCCCAGCAA

714 GTGGTGCCCTACCCTCAGAGAGCTGTGCCTGTTCAAGCCCTTCTGCTCAACCAAGAAC

715 TTCTACTTAACCCCACCCACCAGATCTACCCTGTGACTCAGCCACTTGCCCCAGTTCAT

716 AACCCCATTAGTGTCAAGGAGGGCAAGGGGAGTCTTCTTAACATGCGGGGACGTGGA

717 GGGAAAATCCCGGCCCCGGATCCATGAAACTTGTCTTCCTCGTCCCGCTGTTGCTCGG

718 GGCCCTCGGACTGTGTCTGGCTGGCCGTAGGAGGAGTGTTCAGTGGTGCGCCGTATCC

719 CAACCCGAGGCCACAAAATGCTTCCAATGGGAAAGGAATATGAGAAAAGTGCGTGGC

720 CCTCCTGTGAGCTGCATAAAGAGAGACTCCCCCATCCAGTGTATCCAGGCCATTGCGG

721 AAAACAGGGCCGATGCTGTGACCCTTGATGGTGGTTTCATATACGAGGCAGGCCTGGC

722 CCCCTACAAACTGCGACCTGTAGCGGTGGAAGTCTACGGGACCAAAAGACAGCCACG

723 AACTCACTATTATGCCTTGGCTGTGGTGAAGAAGGGTGGCAGCTTTTAGCTGAACGAA

724 CTGCGAGGTCTGAAGTCCTGCCACACAGGCCTTCGCAGGACCGCTGGATGGGATGTCC

725 CTATAGGGACACTTCGTCCATTCTTGAAATGGACGGGTCCACCTGAGCCCATTGAGGC

AGCTGTGGCCAGGTTCTTCTCAGCCAGCTGTGTTCCCGGTGCAGATAAAGGACAGTCC
CCCAACCTGTGTCGCCTGTGTGCGGGGACAGGGGAAAACAAATGTGCCTTCTCCTCCC
AGGAACCGTACTTCAGCTACTCTGGTGCCTTCGAGTGTCTGAGAGACGGGGCTGGAG
ACGTGGCTTTTATCAGAGAGAGCACAGTGTTCGAGGACCTGTCCGACGAGGCTGAAG
GGGACGAGTATGAGTTACTCTGCCCAGACAACACTCGGAAGCCAGTGGACAAGTTCA
AAGACTGCCATCTGGCCCGGGTCCCTTCTCATGCCGTTGTGGCACGAAGTGTGAATGG
CAAGGAGGATGCGATCTGGGATCTTTCTCCGCCAGGCACAGGAAAAGTTTGGAAAGG
ACAAGTCACCAAAATTCCAGCTCTTTGGCTCCCCTAGTGGGCAGAAAGATCTGCTGTT
CAAGGACTCTGCCATTGGGTTTTCGAGGGTGCCCCCGAGGATAGATTCTGGGCTGTAC
CTTGGCTCCGGCTACTTCACTGCCATCCAGAACTTGAGGAAAAGTGAGGAGGAAGTG
GCTGCCCGGTGTGCGCGGGTCGTGTGGTGTGCGGTGGGCGAGCAGGAGCTGCGCAAG
TGTAACCAGTGGAGTGGCTTGAGCGAAGGCAGCGTGACCTGCTCCTCGGTCTCCACCA
CAGAGGACTGCATCGCCCTGGTGCTGAAAGGAGAAGCTGATGCCATGAGTTTGGATGG
AGGATATGTGTACACTGCAGGCAAGATGTGGTTTGGTGCCTGTCTTGGGAGAGAACTA
CAAATCCCAACAAAGCAGTGACCCTGATCCTAACTGTGTGGATAGACCTGTGGGAGGA
TATCTTGCTGTGGCGGTGGTTAGGAGATCAGACACTAGCCTTACCTGGGACTCTGTGAA
GGGCAAGAGGTCCTGCCACACCGCCGTGGACAGGACTGCAGGCTGGGATATCCCCAT
GGGCCTGCTCTTCAACCAGACGGGCTCCTGCAATTTGATGAGTATTTTAGTCAAGACTG
TGCCCCTGGGTCTGACCCGAGATCTAATCTCTGTGCTCTGTGTATTGGCGACGAGCAGG
GTGAGAATAAGTGCGTGCCCAGCAGCAACGAGAGATACTACGGCTACACTGGGGCTTT
CCGGTGCCTGGCTGAGAATGCTGGAGACGTTGCATTTGTGAAAGATGTGACTGTCTTG
CTGGAACACTGATGGAAATAACAATGAGGCATGGGCTAAAGATTTGAAGCTGGCTGAC
TTTGCGCTGCTGTGCCTCGATGGCAAGCGGGAGCCTGTGACTGAGGCTAGAAGCTGCC
ATCTTGCCATGGCCCCGAATCATGCCGTGGTGTCTCGGATGGATAAAGTGGGACGCCTG
AAGCAGGTGTTGCTCCACCAGCAGGCTAAATTTGGGAGAAATGGATCTGACTGCCCCG
ACAAGTTTTGCTTATTCCAGTCTGAAACCAAAAACCTTCTGTTCAATGACAACACTGA
GTGTCTGGCCAGACTCCATGGGAAAACAACATATGAAAAATATTTGGGACCACAGTAT
GTCTCAGGGCATTACTAATCTGAAAAAGTGCTCAACCTCCCCCCTCCTGGGAGCCTGT
GAATTCCTCAGGAAGTAAAAGCTTGAGAGACGGAGTCACTGCCAACCGAGACGGTCA
TAGCTGTTTCCTGTGTGCCGCTTCCTCGCTCACTGACTCGCTGCGCTCGGTCGTTCGGC

TGCGGCGAGCGGTATCAGCTCACTCAAGGCGGTAATACGGTTATCCGGCGCGCCTCCT
CAGGAAGTAAAAGCTTATAACTTCGTATAGCATACATTATACGAGGTTATACTAGTATTAT
GCCCAGTACATGACCTTATGGGACTTTCCTACTTGGCAGTACATCTACGTATTAGTCATC
GCTATTACCATGGTGATGCGGTTTTGGCAGTACATCAATGGGCGTGGATAGCGGTTTGA
CTCACGGGGATTTCCAAGTCTCCACCCCATTGACGTCGATGGGAGTTTGTTTTGGCACC
AAAATCAACGGGACTTTCCAAAATGTCTTAACAACTCCGCCCCATTGACGCAATGGGC
GGTAGGCGTGTACGGTGGGAGGTCTATATAAGCAGAGCTCGTTTAGTGAACCGTCTGAT
CGCCTGGAGACGCCATCCACGCTGTTTTGACCTCCATAGAAGATTTTAGAGCTAGCGA
AATTCGAATTTAAATCGGATCCGCGGCCGCAAGGATCTGCGATCGCTCCGGTGCCCGTC
TGTGGGCAGAGCGCACATCGCCCACAGTCCCCGAGAAGTTGGGGGGAGGGGTCGGCA
ATTGAACGGGTGCCTAGAGAAGGTGGCGCGGGGTAAACTGGGAAAGTGATGTCGTGT
ACTGGCTCCGCCTTTTTCCCGAGGGTGGGGGAGAACCGTATATAAGTGCAGTAGTCGC
CGTGAACGTTCTTTTTCGCAACGGGTTTGCCGCCAGAACACAGCTGAAGCTTCGAGGG
GCTCGCCATCTCTCCTTCACGCGCCCGCCGCCCTACCTGAGGCCGCCATCCACGCCGGT
TGAGTCGCGTTCTGCCGCCTCCCGCTTGTGGTGCCTCCTGAACTGCGTCCGCCGTCTAG
GTAAGTTTAAAGCTCAGGTCGAGACCGGGCCTTTGTCCGGCGCTCCCTTGGAGCCTAC
CTAGACTCAGCCGGCTCTCCACGCTTTGCCTGACCCTGCTTGCTCAACTCTACGTCTTT
GTTTCGTTTTCTGTTCTGCGCCGTTACAGATCCAAGCTGTGACCGGTGCCTACACCATG
GAGAGCGACGAGAGCGGGCTGCCTGCCATGGAGATCGAGTGCCGCATCACCGGGACC
CTGAACGGTGTGGAGTTTGAGCTGGTGGGCGGTGGAGAGGGTACCCCCAAGCAGGGC
CGCATGACCAGCAAGATGAAGAGCACCAAAGGCTCCCTGACCTTCAGCCCCTACCTGC
TGAGCCACGTGATGGGCTACGGCTTCTACCACTTCGGCACCTACCCCAGCGGCTACGA
GAACCCCTTCCTGCACGCCATCAACAACGGTGGCTACACCAACACCCGCATCGAGAAG
TACGAGGACGGTGGTGTGCTGCACGTGAGCTTCAGCTACCGCTACGAGGCCGGCCGC
GTGATCGGTGACTTCAAGGTGGTGGGCACCGGCTTCCCCGAGGACAGCGTGATCTTCA
CCGACAAGATCATCCGCAGCAACGCCACCGTGGAGCACCTGCACCCCATGGGCGATAA
CGTGCTGGTGGGCAGCTTCGCCCGCTCCTTCAGCCTGCGCGACGGTGGCTACTACAGC
TTCGTGGTGGACAGCCACATGCACTTCAAGAGCGCCATCCACCCCAGCATCCTGCAGA
ACGGGGGCCCCATGTTTGCCCTCCGCCGCGTGGAGGAGCTGCACAGCAACACCGAGC
TGGGCATCGTGGAGTACCAGCACGCCTTCAGGACCCCCATCGCCCTCGCCAGATCCCG

CCGTCAGTCGTCCAATTCTGCCTTGGACGGTACCGCCGGACCCGGCTCCACCGGATCT
CGCTCTGGTGGTTCTCCCAGGAAGAAAAGGAAAGTCGAGGGTAGAGGAAGTCTTCTA
ACATGCGGTGACGTGGAGGAGAATCCCGGCCCTATGACCGAGTACAAGCCCACGGTG
CGCCTCGCCACCCGCGACGACGTCCCCAGGGCCGTACGCACCCTCGCCGCCGCGTTCG
CCGACTACCCCGCCACGCGCCACACCGTGGATCCGGACCGCCACATCGAGCGGGTCAC
CGAGCTGCAGGAACTCTTCCTCACGCGCGTCGGGCTCGACATCGGCAAGGTGTGGGT
CGCGGACGACGGTGCCGCGGTGGCGGTCTGGACCACGCCGGAGAGCGTGGAAGCGG
GGGCGGTGTTCGCCGAGATCGGCCCGCTCATGGCCGAGTTGAGCGGTTCCCGGCTGGC
CTCGCAGCAACAGATGGAAGGCCGCCTGGTGCAGCACCGGCCCAAGGAGCCCGCGTG
GTTCCTGGCCACCGTGGGTGTCTCGCCCGACCACCAGGGCAAGGGTCTGGGCAGCGC
CGTTGTGCTCCCCGGAGTGGAGGCGGCCGAGCGCTCCGGGGTGCCCGCCCTCCTGGA
GACCTCCGCGCCCCGCAACCTCCCCCTTTTACGAGCGGCTCGGCTTCACCGTGACCGC
CGACGTGGAGGTGCCCGAGGGACCGCCCACCTGGTGCATGACCCGCAGGCCCGGTGC
CTGAAGTCAACCTCTGGATTACAAAATTTGTGAAAGATTGACTGGTATTCTTAACTATG
TTGCTCCTTTTACGCTATGTGGATACGCTGCTTTTATGCCTTTGTATCATTCTCTGGTGGT
TCTCCCAAGAAGAAAAGGAAAGTCTAAAACTTGTTTATTGCAGCTTATAATGGTTACAA
ATAAAGCAATAGCATCACAAATTTTACAAATAAAGCATTTTTTTCACTGCATTCTAGTTG
TGGTTTGTCCAAACTCATCAATGTATCTTAAATAACTTCGTATAGCATACATTATACGAG
GTTATATCGATATTGTGGAAAGCCTTTCAAGCAGTGAGGTAAGATAGTGTTTCATTCAG
AGGCATTTTCCCAAATTTAGAGCAATAAAATGCTGTATTATCTTTTTGTGTTACATTAATG
GCAACCCACTCCAGTATTCTTGCCTGGAAGATCCCAAGAGGAGGAGCCTGGTAGGCTG
CAGTCCACGGGGTCGCTAAGAGTCGGACAGGACTGAGCGACTTCCTTTTCACTTTTCA
CTTTTATGCCTTGGAGAAGGAAATGGCAACCCACTCCAGTTTTCTTGCCGGGAGAATC
CCTAGGACGGGGGAGCCTGGTGGGCTGCCTGCTATGGGGTCGCACAGAGTTGGACAC
GACTGAAGCGACTTAGCAGTAGCAGCAGCAGGACAGTTAAGGTTTCTCTAATAGCTCA
GTTGGTAAAGAATCTGCCTGCAGTGCAGGAGACCCTGGTTCGATTGCTCAAGATCTGC
AGGAGAAGGGATAGGCTACCCATTCCAGTGTTCTTTAAAGCCAATGTGGCTATGTACTG
ACGGGTAACTATTGTCAATTTCCACTCTGTATATTTAAGGAATAAATGTGTAGAAGGTTT
AATATTCTAGTAATTTCTAAATGGGTTTGTTATTTGAAATTGTGTCATTGTGCCTTGCTTT
TTTTCCTTAATGAACTGTACAGTCCTCTTTTCTGTTCTTGAGCTTTCTATGTTTACCTCCT

TTTATGCTTTGGCATTTATTGAGCGCTTTCTGTTTTCAGACTTTGACTAGGAACTGCAGT
ACAAACTAGAAAGAGGGATGCCCTTTGTAAGGTGTGAGCAGACATGCAATGGACATAT
TTTATTATTATACAAGCAATCCAGTACACAAGAGGCAGTGAGAATGAGTGTAGTCCTAA
ATCTGCCTGGTGGGATGAGGTAGATAATAACCCTATGCCACTCTTTTTGGCTCTGTGATC
AGCTGTTGGTTAAATAGTGCTTCATATACTTTGTCTCTTCACAAAGGTGAAAAGATGCT
GCTGCTGCTGCTAAGTCGCTTCAGTCATGTCGGACTCTGTGCGACCTCATAGATGGTAG
CCTGCCAGGCTCCCCCATCCCTGGGATTCTCCAGGCGAGAATACTGGAGTGGGTTGCC
ATTTTCTATTTCCATTGCATAAAAGTGAAGAGTGAAGGTGAAGTCGCTCAGTCGTGTCC
GACTCTTAGTGACCCCGTGGACTGCAGCCTACCAGGCTCCTCCATCCATGGGATTTTCC
AGGCGAGGATACTGGAGTGGGTTGCCATTGCCATCTCCAAAGATCCTTTTAGGAGGGT
ATGTATTTCTTATAATTCATTAGAAGCCTAAACATAACCAGGGGACTGAGGATGATTAAC
AAGTTCATGCGTGGTTATTATTATATATTTTCAATGACTATTATTCTTTTAGAACCAGATG
AAAATATTAGAGATCATTTGTTGTCTTAGGGAGAGAACAGGATGATTGAGAGACATGTA
TGCGTGCAAGTTACTTCAGCCCTTTCCAACTCTGTATGACCCTATGGACTGTAACCTGC
CAGGTTTCTCTGTCTATGGGATTCTCCAGGCAAGAATACAGGAGTGGGTTGTCATCTCC
TCCAAGGCCTAGGAATTGACCTGCATCTCTTATGTCTCCTGCATTGGTAGGCTGGTTCTT
TACCACTAGCACCACCTGGGAAGCCTGATTAGTATCTGTTAAATGCCTCTTTGAGTACT
ATGCTCTCTAATCCTCTTTTTCTTGATTGCATCATCTTTCTTTTTATACGACAGCCTTATTC
AGAAGAGTGGGACATAAACTTTTAGCCATAAAATAATGATATTATCGAAAGAGCTGTCC
ATATTAATCTATTAAATTTCTTCATTTTCTGATTTATGTTGACAAATAAGAATTTTTTTAA
AGCTAGACCTGATTTTATTTTTATTTTTCCAAAGGAATCTATTACACGCTTCAATAAGGT
AAAACCCTCATATTTAAATGTACATTTTTTTAAATTTTATGTTTGATTTTTATAAACAGCA
TTTCTTTATGTATTTTTTTTTTAACCAGAAAATTGAGAAGTTTCAGAGTGAGGAACAGC
AGCAAGCAGAGTTTATTAGGCAAAAGGCCAGCCAAAGGCCAGGAACCGTAAAAAGGC
CGCGTTGCTGGCGTTTTTCCATAGGCTCCGCCCCCCTGACGAGCATCACAAAAATCGA
CGCTCAAGTCAGAGGTGGCGAAACCCGACAGGACTATAAAGATACCAGGCGTTTCCCC
CTGGAAGCTCCCTCGTGCGCTCTCCTGTTCCGACCCTGCCGCTTACCGGATACCTGTCC
GCCTTTCTCCCTTCGGGAAGCGTGGCGCTTTCTCATAGCTCACGCTGTAGGTATCTCAG
TTCGGTGTAGGTGGTTCGCTCCAAGCTGGGCTGTGTGCACGAACCCCCCGTTCAGCCC
GACCGCTGCGCCTTATCCGGTAACTATCGTCTTGAGTCCAACCCGGTAAGACACGACTT

ATCGCCACTGGCAGCAGCCACTGGTAACAGGATTAGCAGAGCGAGGTATGTAGGCGGT
GCTACAGAGTTCTTGAAATGGTGGCCTAACTACGGCTACACTAGAAGAACAGTATTTG
GTATCTGCGCTCTGCTGAAGCCAGTTACCTTCGGAAAAAGAGTTGGTAGCTCTTGATCC
GGCAAACAAACCACCGCTGGTAGCGGTGGTTTTTTTGTTTGCAAGCAGCAGATTACGC
GCAGAAAAAAAGGATCTCAAGAAGATCCTTTGATCTTTTCTACGGGGTCTGACGCTCA
GTGGAACGAAAACTTACGTTTAGGGATTTTGGTCATGAGATTATCAAAAAGGATCTTCA
CCTAGATCCTTTTAAATTAAAAATGAAGTTTTAAATCAATCTAAAGTATATATGAGTAAA
CTTGGTCTGACAGTTACCAATGCTTAATCAGTGAGGCACCTATCTCAGCGATCTGTCTAT
TTGGTTATCCATAGTTGCCTGACTCCCCGTCGTGTAGATAACTACGATACGGGAGGGCT
TACCATCTGGCCCCAGTGCTGCATGATACCGCGGGACCCACGCTCACCGGCTCCAGA
TTTATCAGCAATAAACCAGCCAGCCGGAAGGGCCGAGCGCAGAAGTGGTCCTGCAGC
TTTATCCGCCTCCATCCAGTCTATTAATTGTTGCCGGGAAGCTAGAGTAAGTAGTTCGCC
AGTTAATAGTTTGCGCAACGTTGTTGCCATTGCTACAGGCTTCGTGGTATCACGCTCGT
CGTTTGGTATGGCTTCATTCAGCTCCGGTTCCCATCGATCAAGGCGAGTTACATGATCC
CCCATGTTGTGCAAAAAAGCGGTTAGCTCCTTCGGTCCTCCGATCGTTGTCAGAAGTA
AGTTGGCCGCAGTGTATCGCTTCATGGTTATGGTAGCACTGCATAATTCTCTTACTGTCA
TGCCATCCGTAAGATGCTTTTCTGTGACTGGTGAGTACTCAACCAAGTCATTCTGAGAA
TAGTGTATGCGGCGACCGAGTTGCTCTTGCCCGGCGTCAATACGGGATAATACCGCGCC
ACATAGCAGAACTTTAAAAGTGCTCATCATTGGAAAACGTTCTTCGGGGCGAAAACTC
TCAAGGATCTTACCGCTGTTGAGATCCAGTTCGATGTAACCCACTCGTGCACCCAACTG
ATCTTCAGCATCTTTTACTTTCACCAGCGTTTCTGGGTGAGCAAAAACAGGAAGGCAA
AATGCCGCAAAAAAGGGAATAAGGGCGACACGGAAATGTTGAATACTCATACTCTTCC
TTTTTCAATATTATTGAAGCATTTATCAGGGTTATTGTCTCATGAGCGGATACATATTTGA
ATGTATTTAG

**SEQ ID No.2：**

AGGCTTTCCACAATCTATAA

**SEQ ID No.3：**

CTGGAAAGAACTCAAATGTACCAGGCTTTCCACAATCTATAA

**SEQ ID No.4:**

AGTAACAGTCTCTAATGATC

**SEQ ID No.5:**

AGAACTCAAATGTACCTGGTG

**SEQ ID No.6:**

GTCGACATTGATTATTGACTAGTTATTAATAGTAATCAATTACGGGGTCATTAGTTCATAG
CCCATATATGGAGTTCCGCGTTACATAACTTACGGTAAATGGCCCGCCTGGCTGACCGC
CCAACGACCCCCGCCCATTGACGTCAATAATGACGTATGTTCCCATAGTAACGCCAATA
GGGACTTTCCATTGACGTCAATGGGTGGACTATTTACGGTAAACTGCCCACTTGGCAGT
ACATCAAGTGTATCATATGCCAAGTACGCCCCCTATTGACGTCAATGACGGTAAATGGC
CCGCCTGGCATTATGCCCAGTACATGACCTTATGGGACTTTCCTACTTGGCAGTACATCT
ACGTATTAGTCATCGCTATTACCATGGGTCGAGGTGAGCCCCACGTTCTGCTTCACTCT
CCCCATCTCCCCCCCCTCCCCACCCCCAATTTTGTATTTATTTATTTTTTAATTATTTTGTG
CAGCGATGGGGGCGGGGGGGGGGGGGGCGCGCGCCAGGCGGGGCGGGGCGGGGCG
AGGGGCGGGGCGGGGCGAGGCGGAGAGGTGCGGCGGCAGCCAATCAGAGCGGCGCG
CTCCGAAAGTTTCCTTTTATGGCGAGGCGGCGGCGGCGGCGGCCCTATAAAAAGCGAA
GCGCGCGGCGGGCGGGAGTCGCTGCGTTGCCTTCGCCCCGTGCCCCGCTCCGCGCCG
CCTCGCGCCGCCCGCCCCGGCTCTGACTGACCGCGTTACTCCCACAGGTGAGCGGGCG
GGACGGCCCTTCTCCTCCGGGCTGTAATTAGCGCTTGGTTTAATGACGGCTCGTTTCTT
TTCTGTGGCTGCGTGAAAGCCTTAAAGGGCTCCGGGAGGGCCCTTTGTGCGGGGGGG
AGCGGCTCGGGGGGTGCGTGCGTGTGTGTGTGCGTGGGGAGCGCCGCGTGCGGCCCG
CGCTGCCCGGCGGCTGTGAGCGCTGCGGGCGCGGCGCGGGGCTTTGTGCGCTCCGCG
TGTGCGCGAGGGGAGCGCGGCCGGGGGCGGTGCCCCGCGGTGCGGGGGGGCTGCGA
GGGGAACAAAGGCTGCGTGCGGGGTGTGTGCGTGGGGGGGTGAGCAGGGGGTGTGG
GCGCGGCGGTCGGGCTGTAACCCCCCCCTGCACCCCCCTCCCCGAGTTGCTGAGCACG
GCCCGGCTTCGGGTGCGGGGCTCCGTGCGGGGCGTGGCGCGGGGCTCGCCGTGCCGG
GCGGGGGGTGGCGGCAGGTGGGGGTGCCGGGCGGGGCGGGGCCGCCTCGGGCCGGG

GAGGGCTCGGGGGAGGGGCGCGGCGGCCCCCGGAGCGCCGGCGGCTGTCGAGGCGC
GGCGAGCCGCAGCCATTGCCTTTTATGGTAATCGTGCGAGAGGGCGCAGGGACTTCCT
TTGTCCCAAATCTGTGCGGAGCCGAAATCTGGGAGGCGCCGCCGCACCCCCTCTAGCG
GGCGCGGGGCGAAGCGGTGCGGCGCCGGCAGGAAGGAAATGGGCGGGGAGGGCCTT
CGTGCGTCGCCGCGCCGCCGTCCCCTTCTCCCTCTCCAGCCTCGGGGCTGTCCGCGGG
GGGACGGCTGCCTTCGGGGGGGACGGGGCAGGGCGGGGTTCGGCTTCTGGCGTGTGA
CCGGCGGCTCTAGAGCCTCTGCTAACCATGTTCATGCCTTCTTCTTTTTCCTACAGCTCC
TGGGCAACGTGCTGGTTATTGTGCTGTCTCATCATTTTGGCAAAGAATTCTGCAGTCGA
CGGTACCGCGGGCCCGGGATCCACCGGTCCCCTCTCCCTCCCCCCCCCCTAACGTTACT
GGCCGAAGCCGCTTGGAATAAGGCCGGTGTGCGTTTGTCTATATGTTATTTTCCACCATA
TTGCCGTCTTTTGGCAATGTGAGGGCCCGGAAACCTGGCCCTGTCTTCTTGACGAGCA
TTCCTAGGGGTCTTTCCCCTCTCGCCAAAGGAATGCAAGGTCTGTTGAATGTCGTGAA
GGAAGCAGTTCCTCTGGAAGCTTCTTGAAGACAAACAACGTCTGTAGCGACCCTTTGC
AGGCAGCGGAACCCCCCACCTGGCGACAGGTGCCTCTGCGGCCAAAAGCCACGTGTA
TAAGATACACCTGCAAAGGCGGCACAACCCCAGTGCCACGTTGTGAGTTGGATAGTTG
TGGAAAGAGTCAAATGGCTCTCCTCAAGCGTATTCAACAAGGGGCTGAAGGATGCCCA
GAAGGTACCCCATTGTATGGGATCTGATCTGGGGCCTCGGTGCACATGCTTTACATGTG
TTTAGTCGAGGTTAAAAAAACGTCTAGGCCCCCCGAACCACGGGGACGTGGTTTTCCT
TTGAAAAACACGATGATAAGCTTGCCACAACCCACAAGGAGACGACCTTCCATGACCG
AGTACAAGCCCACGGTGCGCCTCGCCACCCGCGACGACGTCCCCCGGGCCGTACGCA
CCCTCGCCGCCGCGTTCGCCGACTACCCCGCCACGCGCCACACCGTCGACCCGGACCG
CCACATCGAGCGGGTCACCGAGCTGCAAGAACTCTTCCTCACGCGCGTCGGGCTCGA
CATCGGCAAGGTGTGGGTCGCGGACGACGGCGCCGCGGTGGCGGTCTGGACCACGCC
GGAGAGCGTCGAAGCGGGGGCGGTGTTCGCCGAGATCGGCCCGCGCATGGCCGAGTT
GAGCGGTTCCCGGCTGGCCGCGCAGCAACAGATGGAAGGCCTCCTGGCGCCGCACCG
GCCCAAGGAGCCCGCGTGGTTCCTGGCCACCGTCGGCGTCTCGCCCGACCACCAGGG
CAAGGGTCTGGGCAGCGCCGTCGTGCTCCCCGGAGTGGAGGCGGCCGAGCGCGCCGG
GGTGCCCGCCTTCCTGGAGACCTCCGCGCCCCGCAACCTCCCCTTCTACGAGCGGCTC
GGCTTCACCGTCACCGCCGACGTCGAGGTGCCCGAAGGACCGCGCACCTGGTGCATG
ACCCGCAAGCCCGGTGCCTAGGCGGCCGCACTCCTCAGGTGCAGGCTGCCTATCAGAA

GGTGGTGGCTGGTGTGGCCAATGCCCTGGCTCACAAATACCACTGAGATCTTTTTCCCT
CTGCCAAAAATTATGGGGACATCATGAAGCCCCTTGAGCATCTGACTTCTGGCTAATAA
AGGAAATTTATTTTCATTGCAATAGTGTGTTGGAATTTTTTGTGTCTCTCACTCGGAAGG
ACATATGGGAGGGCAAATCATTTAAAACATCAGAATGAGTATTTGGTTTAGAGTTTGGC
AACATATGCCCATATGCTGGCTGCCATGAACAAAGGTGGCTATAAAGAGGTCATCAGTA
TATGAAACAGCCCCCTGCTGTCCATTCCTTATTCCATAGAAAAGCCTTGACTTGAGGTT
AGATTTTTTTTATATTTTGTTTTGTGTTATTTTTTTCTTTAACATCCCTAAAATTTTCCTTA
CATGTTTTACTAGCCAGATTTTTCCTCCTCTCCTGACTACTCCCAGTCATAGCTGTCCCT
CTTCTCTTATGAAGATCCCTCGACCTGCAGCCCAAGCTTGGCGTAATCATGGTCATAGC
TGTTTCCTGTGTGAAATTGTTATCCGCTCACAATTCCACACAACATACGAGCCGGAAGC
ATAAAGTGTAAAGCCTGGGGTGCCTAATGAGTGAGCTAACTCACATTAATTGCGTTGCG
CTCACTGCCCGCTTTCCAGTCGGGAAACCTGTCGTGCCAGCGGATCCGCATCTCAATTA
GTCAGCAACCATAGTCCCGCCCCTAACTCCGCCCATCCCGCCCCTAACTCCGCCCAGTT
CCGCCCATTCTCCGCCCCATGGCTGACTAATTTTTTTTATTTATGCAGAGGCCGAGGCCG
CCTCGGCCTCTGAGCTATTCCAGAAGTAGTGAGGAGGCTTTTTTGGAGGCCTAGGCTTT
TGCAAAAAGCTAACTTGTTTATTGCAGCTTATAATGGTTACAAATAAAGCAATAGCATC
ACAAATTTCACAAATAAAGCATTTTTTTCACTGCATTCTAGTTGTGGTTTGTCCAAACTC
ATCAATGTATCTTATCATGTCTGGATCCGCTGCATTAATGAATCGGCCAACGCGCGGGGA
GAGGCGGTTTGCGTATTGGGCGCTCTTCCGCTTCCTCGCTCACTGACTCGCTGCGCTCG
GTCGTTCGGCTGCGGCGAGCGGTATCAGCTCACTCAAAGGCGGTAATACGGTTATCCA
CAGAATCAGGGGATAACGCAGGAAAGAACATGTGAGCAAAAGGCCAGCAAAAGGCC
AGGAACCGTAAAAAGGCCGCGTTGCTGGCGTTTTTCCATAGGCTCCGCCCCCCTGACG
AGCATCACAAAAATCGACGCTCAAGTCAGAGGTGGCGAAACCCGACAGGACTATAAA
GATACCAGGCGTTTCCCCCTGGAAGCTCCCTCGTGCGCTCTCCTGTTCCGACCCTGCCG
CTTACCGGATACCTGTCCGCCTTTCTCCCTTCGGGAAGCGTGGCGCTTTCTCAATGCTC
ACGCTGTAGGTATCTCAGTTCGGTGTAGGTCGTTCGCTCCAAGCTGGGCTGTGTGCAC
GAACCCCCCGTTCAGCCCGACCGCTGCGCCTTATCCGGTAACTATCGTCTTGAGTCCAA
CCCGGTAAGACACGACTTATCGCCACTGGCAGCAGCCACTGGTAACAGGATTAGCAGA
GCGAGGTATGTAGGCGGTGCTACAGAGTTCTTGAAGTGGTGGCCTAACTACGGCTACA
CTAGAAGGACAGTATTTGGTATCTGCGCTCTGCTGAAGCCAGTTACCTTCGGAAAAAG

AGTTGGTAGCTCTTGATCCGGCAAACAAACCACCGCTGGTAGCGGTGGTTTTTTTGTTT
GCAAGCAGCAGATTACGCGCAGAAAAAAAGGATCTCAAGAAGATCCTTTGATCTTTTC
TACGGGGTCTGACGCTCAGTGGAACGAAAACTCACGTTAAGGGATTTTGGTCATGAGA
TTATCAAAAAGGATCTTCACCTAGATCCTTTTAAATTAAAAATGAAGTTTTAAATCAATC
TAAAGTATATATGAGTAAACTTGGTCTGACAGTTACCAATGCTTAATCAGTGAGGCACC
TATCTCAGCGATCTGTCTATTTCGTTCATCCATAGTTGCCTGACTCCCCGTCGTGTAGAT
AACTACGATACGGGAGGGCTTACCATCTGGCCCCAGTGCTGCAATGATACCGCGAGAC
CCACGCTCACCGGCTCCAGATTTATCAGCAATAAACCAGCCAGCCGGAAGGGCCGAGC
GCAGAAGTGGTCCTGCAACTTTATCCGCCTCCATCCAGTCTATTAATTGTTGCCGGGAA
GCTAGAGTAAGTAGTTCGCCAGTTAATAGTTTGCGCAACGTTGTTGCCATTGCTACAGG
CATCGTGGTGTCACGCTCGTCGTTTGGTATGGCTTCATTCAGCTCCGGTTCCCAACGAT
CAAGGCGAGTTACATGATCCCCCATGTTGTGCAAAAAAGCGGTTAGCTCCTTCGGTCC
TCCGATCGTTGTCAGAAGTAAGTTGGCCGCAGTGTTATCACTCATGGTTATGGCAGCAC
TGCATAATTCTCTTACTGTCATGCCATCCGTAAGATGCTTTTCTGTGACTGGTGAGTACT
CAACCAAGTCATTCTGAGAATAGTGTATGCGGCGACCGAGTTGCTCTTGCCCGGCGTC
AATACGGGATAATACCGCGCCACATAGCAGAACTTTAAAAGTGCTCATCATTGGAAAAC
GTTCTTCGGGGCGAAAACTCTCAAGGATCTTACCGCTGTTGAGATCCAGTTCGATGTAA
CCCACTCGTGCACCCAACTGATCTTCAGCATCTTTTACTTTCACCAGCGTTTCTGGGTG
AGCAAAAACAGGAAGGCAAAATGCCGCAAAAAAGGGAATAAGGGCGACACGGAAAT
GTTGAATACTCATACTCTTCCTTTTTCAATATTATTGAAGCATTTATCAGGGTTATTGTCT
CATGAGCGGATACATATTTGAATGTATTTAGAAAAATAAACAAATAGGGGTTCCGCGCA
CATTTCCCCGAAAAGTGCCACCTG
